# USP8 Controls Proteostasis Pathways in B Cells and Multiple Myeloma

**DOI:** 10.1101/2024.04.26.591134

**Authors:** Almut Dufner, Fabien Thery, Gianni Monaco, Jelena Lazarevic, Oliver Gorka, Nina Chevalier, Marta Campos Alonso, Maximilian Frosch, Gerbrand J. van der Heden van Noort, Kira Allmeroth, Marco Prinz, Olaf Groß, Huib Ovaa, Paul P. Geurink, Wolfgang W. Schamel, Martin S. Denzel, Vigo Heissmeyer, Benedikt Jacobs, Mirle Schemionek, Francis Impens, Heiko Bruns, Klaus-Peter Knobeloch

## Abstract

Ubiquitin-specific protease 8 (USP8) is a multifunctional deubiquitinating enzyme that plays a pivotal role in the regulation of endosomal and lysosomal trafficking. Several studies showed that USP8 is critically involved in the pathogenesis of various tumor entities.

Recently, USP8 emerged as a vulnerability gene in multiple myeloma (MM), suggesting a functional role in the B- and plasma cell compartment. Here we have comprehensively analyzed mice with deletion of *Usp8* at different stages of B-cell development. Furthermore, we evaluated the function of USP8 in proteasome inhibition susceptible and Bortezomib (BTZ) resistant patient derived MM cells using depletion of USP8 and treatment with DUB-IN-2, a widely used inhibitor published to target USP8.

*Usp8* depletion in *Usp8*^f/f^C*d19*-Cre mice affected B-cell survival and development favoring immature, innate-like B cells, and germinal center and plasma cells, while also elevating immune-responses and causing Roquin depletion. Cells expressing catalytically inactive USP8 accumulated proteins modified with mixed ubiquitin/NEDD8 chains, indicating proteotoxic stress. Moreover, we identified these modifications as preferred substrates of USP8. In MM cells, efficient USP8 knockdown reduced survival by inducing lysosomal dysfunction. In contrast, DUB-IN-2 induced an enhanced ER stress response to treatment with BTZ.

Our results highlight the potential of targeting USP8 and the combination of DUB-IN-2 and BTZ in treating BTZ-resistant MM. However, our biochemical and cellular analyses raise fundamental concerns about the function of DUB-IN-2 as a USP8 inhibitor.

## Introduction

Ubiquitin modification represents a highly versatile mechanism to control cellular functions. Targeting the ubiquitin proteasome system (UPS) represents an effective strategy for the treatment of B-cell malignancies such as multiple myeloma (MM)[1]. However, although proteasome inhibition by bortezomib (BTZ, Velcade®) or Carfilzomib (CFZ, Kyprolis®) enhanced patient outcome, relapse is common in MM and limits therapeutic efficacy. Ubiquitylation is counteracted by the activity of ubiquitin deconjugating enzymes (DUBs)[2]. More than 90 different DUBs including around 50 members of the ubiquitin-specific protease (USPs) family exist. USP8 is characterized by a multi-domain architecture[3]. Interactions with components of endosomal sorting complexes required for transport (ESCRT), such as STAM2 and charged multivesicular body (CHMP) proteins, have established the regulation of endosomal sorting and transmembrane protein stability as canonical USP8 functions[3]. The C-terminal catalytic domain of USP8 is inhibited by phosphorylation-dependent 14-3-3 binding to the adjacent 14-3-3 binding motif (BM)[4,5]. Somatic mutations affecting the 14-3-3 BM are associated with Cushing’s disease (CD) leading to EGFR stabilization in ACTH-secreting pituitary adenomas[6,7]. USP8 was also associated with autophagy and mitophagy regulation[3,8,9], and with the progression of many types of cancer including effects on the tumor microenvironment via regulation of PD-L1, CTLA-4, TGF-β signaling, and inflammatory responses[10–14]. Numerous studies investigating the role of USP8 in cancer utilized DUB-IN-2, a previously reported USP8 inhibitor [10–12,15–17]. However, the mechanism of action of this compound remains unclear and specificity and kinetic parameters have not been investigated in detail[18,19].

In T cells, USP8 is essential for the positive selection of thymocytes[20]. Mice with a T-cell-specific deletion of USP8 have a defect in homeostatic and antigen-driven T-cell expansion. They spontaneously develop colitis concomitant with dysfunctional regulatory T cells. Moreover, USP8 interacts with GRB2-related adaptor protein downstream of Shc (GADS) and controls stability of the ESCRT component CHMP5 to stabilize the anti-apoptotic Bcl2 protein as a prerequisite for the positive selection of thymocytes[20,21].

Ubiquitylation and deubiquitylation are also important processes in the regulation of B-cell activation, proliferation, and apoptosis during development and their functional diversification and activation in innate and humoral immune responses[22]. A systems-wide analysis of B-cell receptor signalosomes has identified USP8 as an interacting component upon stimulation[23]. Interest in the function of USP8 in the B cell compartment is further fueled by an RNA interference lethality screen, which identified *Usp8* as a key gene exhibiting a marked difference in functional vulnerability between MM and non-MM cells[24]. MM cells, which represent cancerous plasma cells, are engaged in rapid protein synthesis, strongly depend on UPS-mediated protein turnover, and are highly susceptible to perturbations of protein homeostasis. Recent studies have demonstrated that inhibiting USPs such as USP7, USP9X, and USP14 effectively promote apoptosis in MM cells, highlighting their potential as therapeutic targets[25,26]. Although USP8 has been implicated in B-cell signaling and MM cell survival, its comprehensive role in B-cell development, immune function, and MM pathogenesis remains unclear.

Here we have comprehensively analyzed mice with deletion of *Usp8* at different stages of B-cell development and evaluated the function of USP8 in MM cells resistant or susceptible to proteasome inhibition. While *Usp8*^f/f^mb1-Cre B cells exhibit an early developmental block, *Usp8*^f/f^*Cd19*-Cre mice are characterized by developmental defects associated with a predominance of immature and innate-like B cells, and germinal center (GCs) and plasma cells (PCs) with elevated immune-responsive functions. Shotgun proteomics and ubiquitinome analyses of B cells, together with signal transduction analyses of both B and MM cells with manipulated USP8 activity, uncovered a fundamental role of USP8 in preventing cell death induced by proteotoxic stress. Of note, USP8 directly controls branched NEDD8/ubiquitin conjugation typically associated with proteotoxic stress. However, while the inhibitor DUB-IN-2, which was reported to target USP8[18], resensitized BTZ resistant MM cells to treatment in a BTZ-synergistic manner through the ER stress response, USP8 knockdown reduced cell survival independently of BTZ by causing lysosomal dysfunction. Our results suggest that targeting USP8 and the combination of DUB-IN-2/BTZ represent promising strategies for overcoming BTZ-resistance in MM. Crucially, our data suggest that DUB-IN-2 does not target USP8, other DUBs or the proteasome, indicating a novel off-target mechanism with therapeutic implications.

## Results

### USP8 is essential for B-cell development

USP8 is critical for T-cell homeostasis where GADS has been identified as a USP8 binding partner[20]. However, B lymphocytes express only marginal levels of GADS (Fig. S1A), suggesting a different mode of action. To evaluate the roles of USP8 in B cells, we generated *Usp8*^f/f^*Cd19*-Cre mice, which delete Usp8 from the pre-B-cell stage on, and compared them to *Usp8*^f/f^ mice (Fig. S1B). USP8 protein expression was not fully abolished in splenic *Usp8*^f/f^*Cd19*-Cre B cells indicating that cells with incomplete Usp8 deletion gain a survival advantage (Fig. S1C).

In contrast to T cells, percentages and total numbers of CD19^+^ B cells are reduced in spleens of *Usp8*^f/f^C*d19*-Cre mice (Fig. 1A). *Usp8*^f/f^*Cd19*-Cre mice displayed a more pronounced reduction of follicular B cells as compared to marginal zone B cells (Fig. 1B), consistent with a defect in the development of mature (IgD^+^) B cells in the spleen (Fig. 1C). Likewise, *Usp8*^f/f^*Cd19*-Cre mice exhibited a significant reduction in the percentages of CD19^+^ and mature IgM^+^IgD^+^ BM B cells, but not early developmental BM B cells (Fig. 1 D). Transitional B-cell development was impaired starting from the T2 stage (Fig. 1E). These results were confirmed using *Usp8*^+/+^*Cd19*-Cre mice as controls (Fig. S1D). Moreover, B1a and B1b subsets, which play key roles in innate immunity, tended to show lower numbers in spleen and peritoneal cavity (PC) in the absence of USP8, while percentages within the IgM^+^ subset increased (Fig. S1E). Analyses of the splenic architecture of *Usp8*^f/f^*Cd19*-Cre mice confirmed the developmental defects of B-cell subsets (Fig. S1F). In *Usp8*^f/f^*Cd19*-Cre spleens, B cell-rich (B220^+^) areas were reduced in size as compared to *Usp8*^f/f^ spleens and in contrast to the T cell-rich periarteriolar lymphoid sheets (CD3^+^). Consistent with the severe reduction of IgD^+^ B cells, the reduction in size of B cell-rich (B220^+^) areas was primarily caused by diminishment of primary B-cell follicles (FO). Moreover, USP8 knockout (KO) cells that have been stimulated by BCR or TLR4/9 engagement exhibit a defect in proliferation (Fig. S1G), and the average number of Ki67 positive cells in splenic B-cell follicles was significantly reduced in *Usp8*^f/f^*Cd19*-Cre mice as compared to controls (Fig. S1H). To determine whether USP8 is critical for earlier steps in B-cell development, we generated *Usp8*^f/f^*mb1*-Cre mice, which delete Usp8 from the early pro B-cell stage on[27]. *Usp8*^f/f^*mb1*-Cre BM B cells showed a block at the pre-B-cell stage as compared to *Usp8*^+/f^*mb1*-Cre B controls (Fig. S1I, upper left panels). As a consequence, a complete loss of peripheral and recirculating B cells was observed (Fig. S1I).

**Fig. 1.**
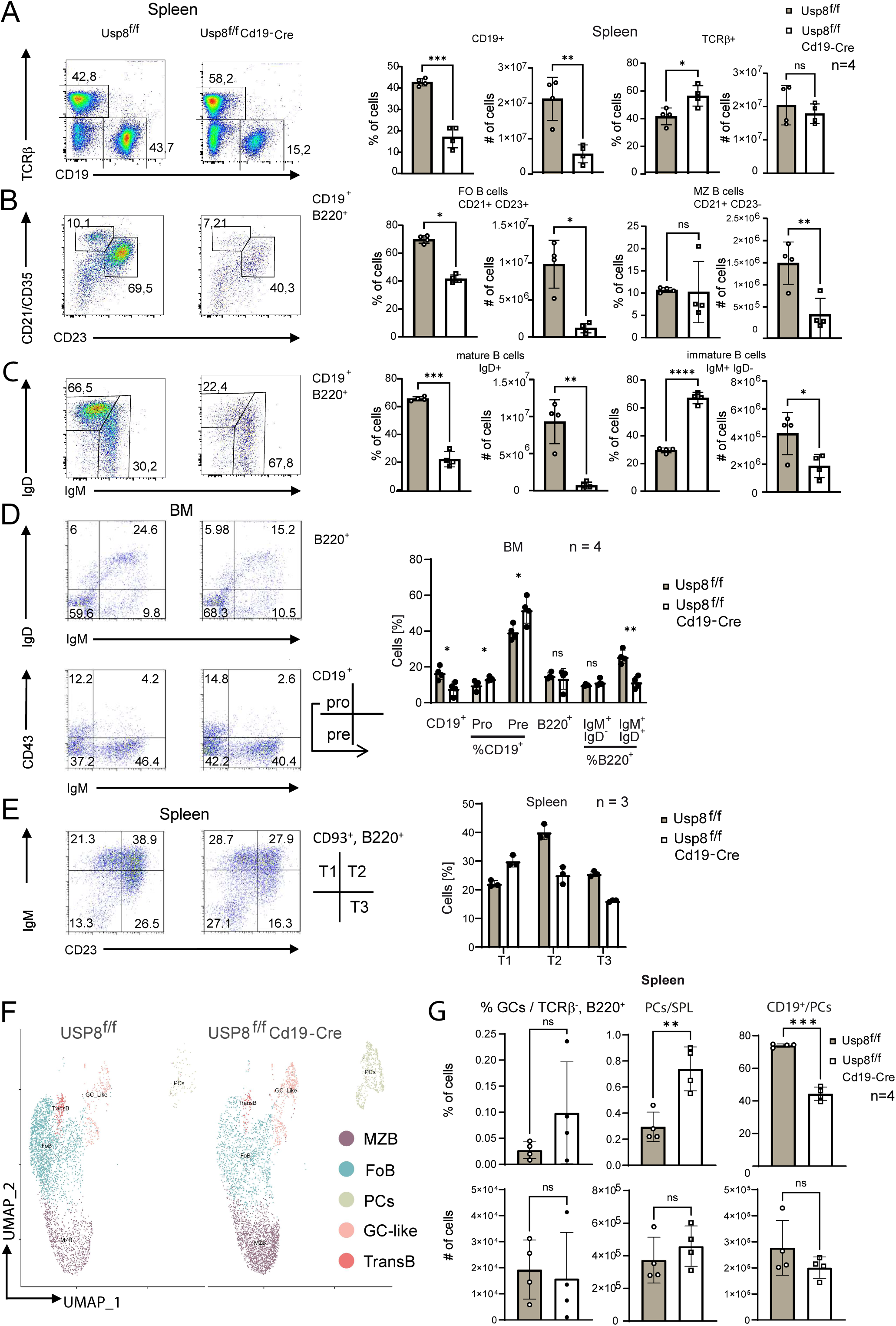
Defect in B-cell development in *Usp8*^f/f^*Cd19*-Cre mice. (A-C) Flow cytometry analysis of B-cell subsets in spleen of *Usp8*^f/f^ and *Usp8*^f/f^*Cd19*-Cre mice (n=4). Pre-gating is indicated at the top right of the FACS plots. Percentages and numbers of B-cell subsets are shown in the right panels. Unpaired 2-tailed t-test, mean+/- sd. (D) Flow cytometry analysis of *Usp8*^f/f^ and *Usp8*^f/f^*Cd19*-Cre bone marrow (BM) (n=4). Paired 2-tailed t-test, mean+/- sd. (E) Flow cytometry analysis of splenic transitional *Usp8*^f/f^ and *Usp8*^f/f^*Cd19*-Cre B-cell subsets (n=3). Paired, 2-tailed t-test, mean+/-sd. (F) Uniform Manifold Approximation and Projection (UMAP) of B cells enriched from spleens of *Usp8*^f/f^ and *Usp8*^f/f^*Cd19*-Cre spleens. (G) Flow cytometry analysis of splenic *Usp8*^f/f^ and *Usp8*^f/f^*Cd19*-Cre germinal center (GC; GL7^hi^, Fas^hi^, TCR-β^−^, B220^+^) B cells and CD138^hi^ PCs, and CD19 expression on PCs (n=4). Unpaired 2-tailed t-test, mean+/- sd. MZB: marginal zone B cells; FoB: follicular B cells; Pro: pro-B cells; Pre: pre-B cells; TransB: Transitional B cells; T1, T2, T3: Transitional T1, T2, T3 B cells; BM: bone marrow; PCs: plasma cells; GC-like: germinal center-like. Paired analysis was used, if sex-matched littermates were derived from n different litters. Source data are provided as source data file.

In summary, our data demonstrate that USP8 is essential for normal B-cell development at various stages.

### Elevated Ig responses and robust PC numbers in *Usp8*^f/f^*Cd19*-Cre mice

To characterize B-cell development in detail, we performed single cell RNA-seq analysis of splenic B cells enriched from *Usp8*^f/f^ and *Usp8*^f/f^*Cd19*-Cre mice. Unbiased clustering of the aggregated dataset produced 10 discrete clusters (Fig. S2A). Although remaining macrophages and granulocytes were overrepresented among enriched *Usp8*^f/f^*Cd19*-Cre B cells, numbers of splenic *Usp8*^f/f^*Cd19*-Cre monocytes/macrophages and granulocytes were not altered significantly (Figure S2B). B cells and PCs were extracted from individual samples and integrated to create a combined dataset. B-cell subpopulations were annotated using top 10 (Fig. S2C) and key signature genes according to Nguyen et al.[28]. As expected, *Usp8*^f/f^*Cd19*-Cre follicular and transitional B cells were diminished as compared to controls (Fig. 1F). Notably, *Usp8*^f/f^*Cd19*-Cre germinal center (GC)-like and PC populations were increased relative to other B-cell subsets as compared to *Usp8*^f/f^ controls (Fig. 1F and 1G upper panels). Total *Usp8*^f/f^*Cd19*-Cre GC B-cell and PC numbers were not significantly affected, including CD19^+^ PCs (Fig. 1G, lower panels). However, CD19^+^ PCs were underrepresented among *Usp8*^f/f^*Cd19*-Cre PCs (Fig. 1G, upper right panel) suggesting that CD19 positive *Usp8*^f/f^*Cd19*-Cre PCs have a survival disadvantage due to more efficient Usp8 deletion. Differential gene expression analysis revealed elevated IL-9R expression in FoB and MZB, elevated expression of specific Ig variable regions in PCs, and reduced expression of ZFP318, a transcriptional regulator of IgD expression[29], in GC-like B cells and FoB of *Usp8*^f/f^*Cd19*-Cre mice (Fig. S2D).

Regarding the functional capabilities of B cells *in vivo*, basal serum levels of all Igs tested did not differ significantly between genotypes (Fig. 2A). Despite a decrease in total GC B-cell numbers in spleens of TNP-KLH-immunized *Usp8*^f/f^*Cd19*-Cre mice, percentages among B cells were not significantly affected (Fig. 2B). We also measured IgM, IgG_1,_ or IgG_3_ responses to defined T cell-dependent (TD) and T cell-independent (TI) immune responses (Fig. 2C). Of note, *Usp8*^f/f^*Cd19*-Cre mice produced significantly elevated levels of IgM and IgG_3_, and spleens of TNP-KLH-immunized *Usp8*^f/f^*Cd19*-Cre mice contained elevated numbers of PCs (Fig. 2D). However, as shown for unimmunized mice, percentages of CD19^+^ PCs within the PC compartment were reduced. Consistent with elevated IgM responses, *Usp8*^f/f^*Cd19*-Cre mice showed higher numbers of IgM-positive, but not IgG1 positive PCs per spleen (Fig. 2E). The expression of Blimp1 and IRF4, two transcription factors essential for PC generation and function[30], remained unchanged (Fig. 2F). Likewise, splenic naïve, effector memory, and regulatory T-cell populations, and basal blood IL1-β, IL-6, and TNF cytokine levels were not affected in *Usp8*^f/f^*Cd19*-Cre mice (Fig. S2E and S2F).

**Fig. 2.**
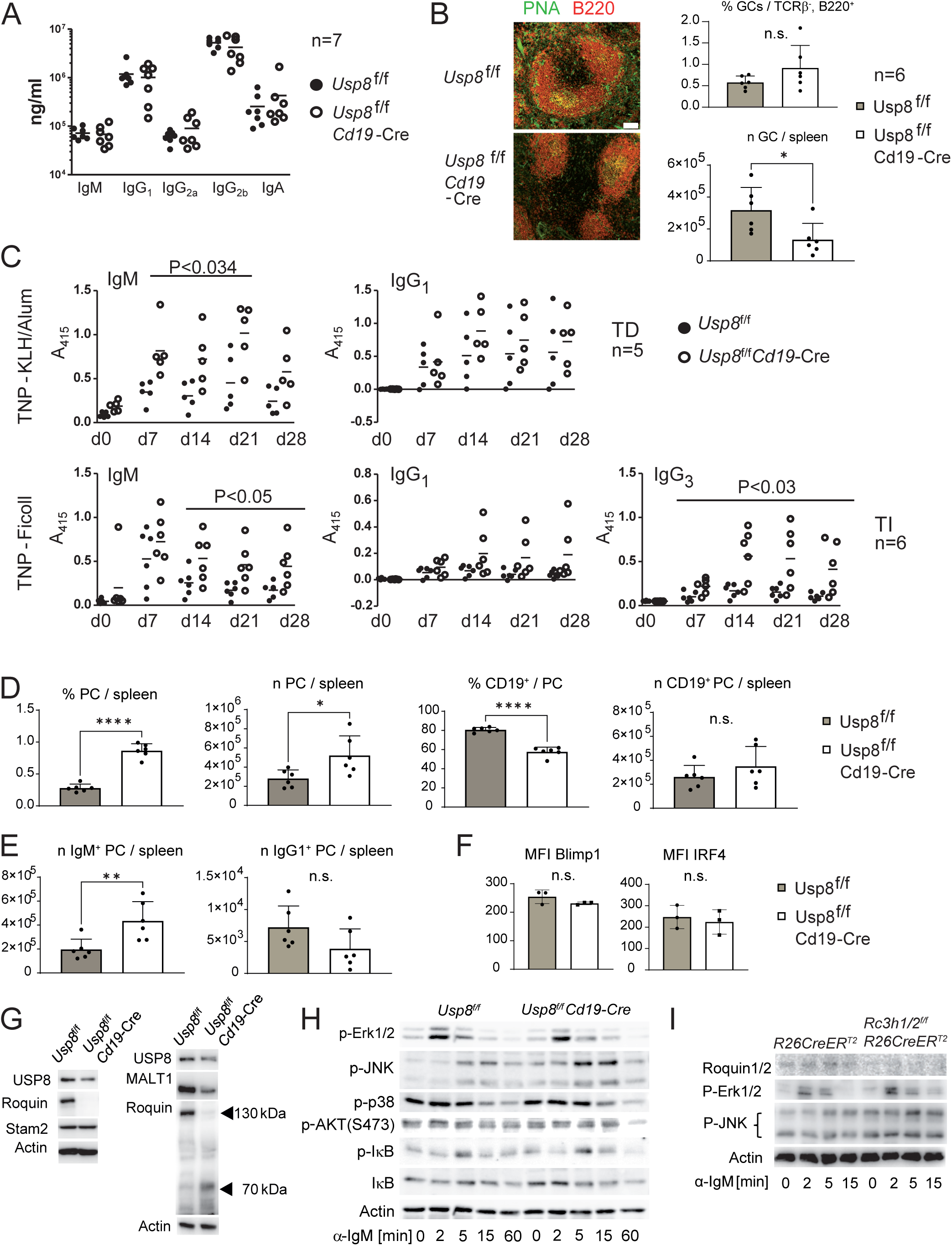
*Usp8*^f/f^*Cd19*-Cre mice exhibit enhanced PC numbers and Ig responses upon T cell-dependent and T cell-independent immunization. (A) Normal basal immunoglobulin titers in *Usp8*^f/f^*Cd19*-Cre mice (n=7). (B) Control and *Usp8*^f/f^*Cd19*-Cre mice were i.p. injected with 50 μg TNP-KLH precipitated in alum and GC B cells were visualized by fluorescence imaging of frozen sections (scale bar, 100 µm) and quantified by FACS analysis (GL7^hi^, Fas^hi^, TCR-β^−^, B220^+^) at day 14 post injection (n=6, unpaired 2-tailed t-test, mean+/-sd). (C) Detection of TNP-specific Ig present in the serum of *Usp8^f/f^*and *Usp8*^f/f^*Cd19*-Cre mice that have been i.p. injected with 50 μg TNP-KLH precipitated in alum (TD response, n=5) or with 25 μg TNP-Ficoll (TI response, n=6) by ELISA. Significant differences are summarized collectively. Unpaired 2-tailed t-test, mean+/-sd. (D-F) *Usp8^f/f^* and *Usp8*^f/f^*Cd19*-Cre mice were treated as in (B) and spleen single cell suspensions were analyzed by FACS to determine frequencies of CD138^hi^ PCs and CD19 expression on PCs (n=6) (D), frequencies of IgM^+^ and IgG1^+^ PCs (n=6) (E), as well as the expression of Blimp1 and IRF4 in PCs (n=3) (F). Unpaired 2-tailed t-test, mean+/-sd. (G) Splenic *Usp8*^f/f^ and *Usp8*^f/f^*Cd19*-Cre B cells were enriched and analyzed by Western blotting (WB). Results are representative of at least 2 independent experiments. (H) Splenic *Usp8*^f/f^ and *Usp8*^f/f^*Cd19*-Cre B cells were starved for 2 hours, stimulated for the indicated times with anti-IgM F(ab’) fragment antibody, and analyzed by WB. Results are representative of at least 2 independent experiments. (I) Upon oral gavage of tamoxifen, Roquin1/2 depleted *Rc3h1/2*^f/f^ *Rosa26-*CreERT2 B cells and *Rosa26-*CreERT2 control cells were enriched from spleens, stimulated as in (H), and analyzed by WB (n=3). Source data are provided as source data file.

Roquin proteins, including Roquin-1 and Roquin-2, are required for shaping T and B cell-dependent immune responses[31,32]. We thus hypothesized that Roquin function might be compromised in *Usp8*^f/f^*Cd19*-Cre B cells. Indeed, Roquin(1/2) appeared to be degraded in splenic USP8-deficient B cells as compared to *Usp8*^f/f^ controls and Stam2, which has been reported to be stabilized by USP8 in other cells, underlining the various functions of USP8 in different cell types[33,34] (Fig. 2G). To evaluate whether USP8 directly controls Roquin stability, we ablated USP8 in *Usp8*^f/f^*Rosa26-*Cre-ERT2 splenic B cells *ex vivo* by administration of 4-hydroxytamoxifen (OHT). Remarkably, short-term *ex vivo* USP8 ablation did not directly affect Roquin protein levels (and did not abolish CHMP5 expression) (Fig. S3A). Thus, altered Roquin protein levels in *Usp8*^f/f^*Cd19*-Cre B cells are rather a consequence of altered B-cell composition and/or signaling within an intact *in vivo* microenvironment rather than a result of the absence of a direct stabilizing effect mediated by USP8 deubiquitylation of Roquin. Roquin cleavage is mediated by Malt1 whose protease function is positively controlled by mono-ubiquitylation at lysine (K) 644 or excess exposure to free ubiquitin[35,36]. Although mono-ubiquitylation of Malt1 was not apparent in *Usp8*^f/f^*Cd19*-Cre B cells (Fig. 2G), gene expression of Usp8, Roquin, and Malt1 correlates accurately across B-cell subtypes (Fig. S3B), suggesting interconnected regulatory mechanisms.

In line with elevated Ig responses, we observed elevated JNK activation (Fig. 2H) and Ca^2+^ mobilization (Fig. S3C) in splenic *Usp8*^f/f^*Cd19*-Cre B-cells in response to anti-IgM-stimulation, while Erk1/2, p38, AKT, and IκB (Fig. 2H) or overall tyrosine phosphorylation (Fig. S3D) was not significantly affected. However, similar to Roquin, *ex vivo* induced depletion of USP8 in *Usp8*^f/f^*Rosa26-*Cre-ERT2 splenic B cells did not significantly affect JNK (Fig. S3E) and Ca^2+^ signaling (Fig. S3F). To assess if elevated JNK activation results from Roquin depletion, we analyzed IgM-induced JNK activation in splenic B cells with *in vivo* Roquin1/2 depletion. Our findings implicate that heightened JNK activation in *Usp8*^f/f^*Cd19*-Cre B cells after anti-IgM stimulation is a direct consequence of *in vivo* Roquin1/2 depletion (Fig. 2I), while Ca^2+^ mobilization remained unaltered (Fig. S3G). Pathway enrichment analysis of different B-cell subsets revealed upregulated BCR second messenger signaling and Rho-GTPase signaling in USP8-deficient FoB cells, as well as upregulated Rho GTPase and death receptor (DR)/TNFR signaling in GC-like USP8-deficient B cells. USP8-deficient MZB cells show upregulation of ER Golgi anterograde transport, p53 signaling, Raf, and CD28 signaling. USP8-deficient PCs vary from control PCs primarily with regard to processing of capped intron-containing mRNA. Of note, also NEDDylation (Table S1) and stress response genes were upregulated in PCs (Fig. S3H). Moreover, elevated rho signaling in diverse *Usp8*^f/f^*Cd19*-Cre B cell subsets may account for the increased Ca^2+^ influx.

In summary, PCs, GC-like B cells, immature, and innate-like B cells dominate the splenic B-cell compartment of *Usp8*^f/f^*Cd19*-Cre mice. USP8-deficient B cells exhibit enhanced Ig, JNK, and Ca^2+^responses, potentially driven by *in vivo* Roquin depletion and altered rho-GTPase signaling[37]. Considering the essential function of USP8, the observed signaling alterations are most likely complex and might also reflect B cell subset-specific interactions with the microenvironment.

### USP8 limits proteotoxic stress and cleaves ubiquitin/NEDD8 mixed chains

To further evaluate the mechanistic role of USP8 in B cell signaling, survival and homeostasis, we stably expressed FLAG-tagged mouse wild-type (WT) USP8 and catalytically inactive mouse USP8(C748A) in the murine K46Lµm B-cell lymphoma line (expressing NP-specific mIgM-BCR)[38,39] (Fig. 3A) and analyzed the proteome and its modification by ubiquitin and ubiquitin-like modifiers using the diGly enrichment method (Ubiscan®)[40]. Overexpression of catalytically inactive variants of USPs is a suitable approach to trap USP substrates[41]. Unexpectedly, NEDD8 levels were elevated in B cells expressing USP8(C748A) (Fig. 3B, left panel). NEDD8 is the ubiquitin-like (ubl) modifier with the highest similarity to ubiquitin[42]. Remarkebly, multiple ubiquitin/ubl modified sites on NEDD8, including K11, K22, K33, K48, K54, and K60, were identified (deposited in PRIDE database, identifier PXD024395; for accession details please see data availability section). These data indicated that free or conjugated NEDD8 levels were increased in the presence of USP8(C748A), and that NEDD8 itself is modified by ubiquitylation or NEDDylation, which cannot be distinguished by the diGly enrichment method. Moreover, enhanced ubiquitin/ubl-modification of specific sites of the ribosomal proteins Rpl7, Rpl8, Rpl11, and ribosome biogenesis regulator 1 (Rrs1) were detected in the presence of USP8(C748A) (Fig. 3B, right panel).

**Fig. 3.**
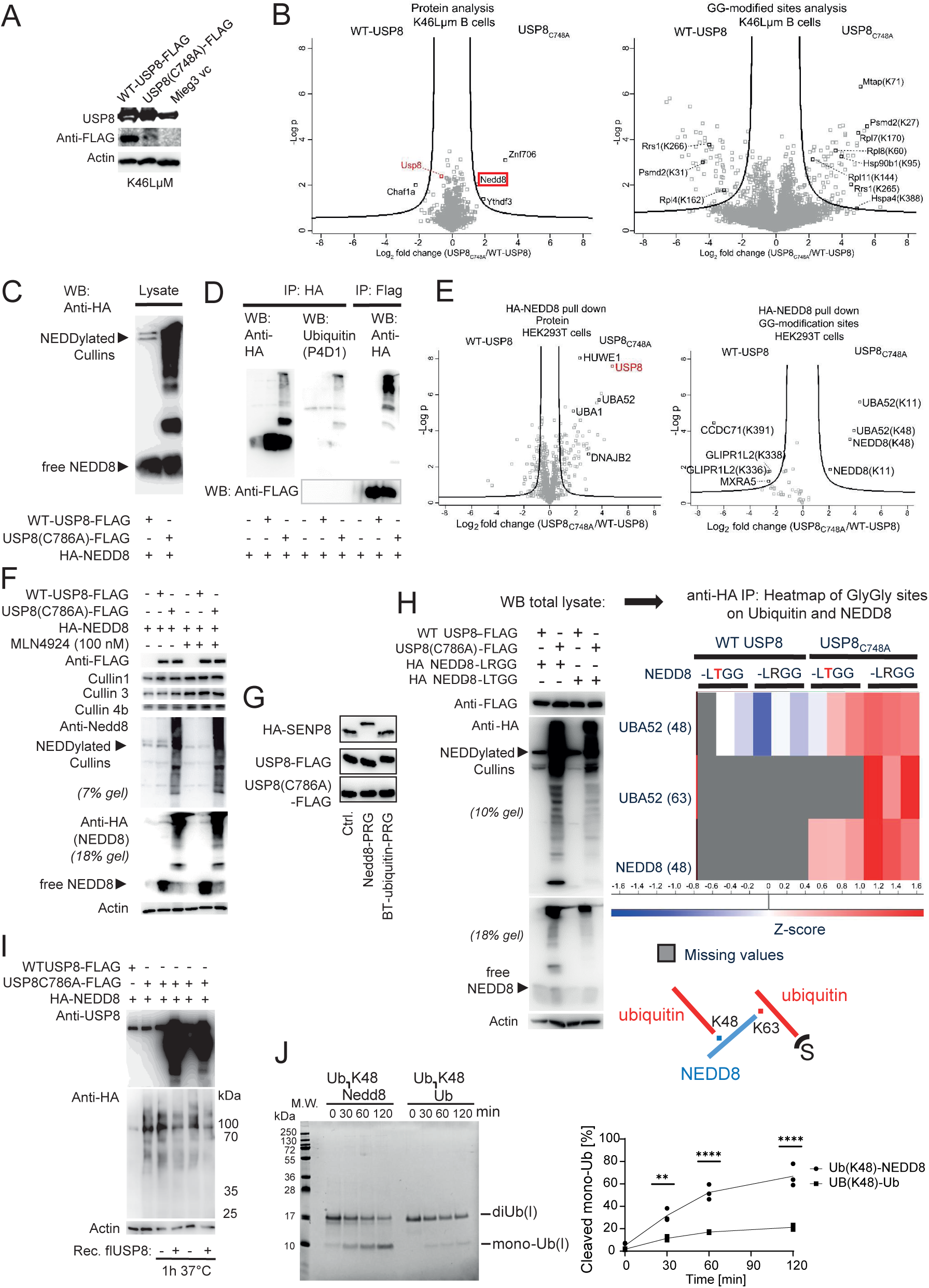
USP8 regulates mixed ubiquitin/NEDD8 chain formation indicating proteotoxic stress. (A, B) NEDD8 levels and ribosomal protein modifications are elevated in B cells expressing USP8(C748A)-FLAG. (A) WT-USP8-FLAG and USP8(C748A)-FLAG expression in K46Lµm B cells analyzed by WB. (B) Volcano plots of USP8(C748A):WT-USP8 diGly proteomics experiments. Differentially expressed proteins (shotgun analysis) and regulated diGly sites (GG-modified sites analysis) were identified in proteins derived from K46Lµm B cells (n=3). Lysine positions of the identified diGly sites are included in parentheses. (C-E) HEK293T cells were transfected with WT-USP8-FLAG or USP8(C786A)-FLAG in the presence of HA-NEDD8. (C) Lysates were analyzed by WB. (D) Anti-HA and anti-FLAG pulldowns were analyzed by WB. (E) Anti-HA pulldowns were analyzed by MS (n=3) to quantify pulled down proteins and GlyGly-K sites. The lysine position of the identified di-glycine site is included in parentheses. (F) HEK293T cells were transfected as in (C). 24 h prior to lysis cells were treated with MLN4924 (100 nM) as indicated and lysates were analyzed by WB. (G) HEK293T cells were transfected with HA-SENP8, WT-USP8-FLAG or USP8(C786A)-FLAG. Lysates were incubated with buffer alone (ctr.), NEDD8-propargylamide (NEDD8-PRG), or ubiquitin-PRG and analyzed by WB using anti-HA or anti-FLAG antibodies. (H) HEK293T cells were transfected with WT-USP8-FLAG or USP8(C786A)-FLAG in the presence of HA-NEDD8(LRGG) or HA-NEDD8(LTGG).Lysates were analyzed by WB (left panels). Anti-HA pulldowns were analyzed by MS (3 replicates/condition) and alterations of ubiquitin (UBA52) and NEDD8 diGly sites are shown in the heat map. Hybrid linkages between both modifiers are reconstructed in the scheme below. (I) Protein lysates generated from 293T cells transfected with USP8(C786A)-FLAG and HA-NEDD8 were incubated with recombinant USP8 as indicated and analyzed by WB. (J) DUB cleavag e assays were performed with recombinant human full length USP8 and synthetic K48-ubiquitinated-NEDD8 (n=3) or K48-diubiquitin (n=2) for the times indicated. The resulting cleavage products were visualized after protein electrophoresis and protein staining of the gel. Statistical analysis after quantification utilized two-tailed unpaired t-test. Source data are provided as source data file.

Similar to ubiquitin conjugation, NEDDylation is a sequential process brought about by the NEDD8 activating enzyme (NAE), a heterodimer of NAE1 and UBA3, the conjugating enzymes UBE2M (UBC12) and UBE2F, and a set of E3 ligases. The best-established NEDDylation targets are cullin proteins, which stimulate cullin-RING E3 ubiquitin ligases (CRLs) upon mono-NEDDylation. In contrast to this canonical NEDDylation, overexpression of NEDD8 or proteasome inhibition leads to a shift in the free ubiquitin-to-NEDD8 ratio which ultimately triggers atypical conjugation of NEDD8 by the ubiquitin activating enzyme E1 (UBE1/UBA1) and the E3 ligase HUWE1 leading to the formation of mixed ubiquitin/NEDD8 chains[43–48] representing a defense mechanism against proteotoxic stress[48]. Ribosomal substrates typically form nuclear aggregates upon modification by ubiquitin/NEDD8 mixed chains to protect the nuclear UPS from an increased load of substrate proteins representing a classic hallmark of proteotoxic stress[48].

Reminiscent of atypical NEDDylation arising as a proteotoxic stress response, overall NEDDylation was strongly enhanced if we co-expressed HA-tagged NEDD8 with human USP8(C786A)-FLAG in HEK293T cells as compared to co-expression with WT USP8 (Fig. 3C). Ubiquitinated proteins were present in the HA-NEDD8 pulldown of these protein lysates, strongly suggesting the presence of mixed ubiquitin/NEDD8 chains (Fig 3D). Vice versa, USP8(C786A)-FLAG pulldowns comprised NEDDylated proteins, but not free NEDD8, indicating that catalytically inactive USP8 traps NEDDylated substrates possibly via ubiquitin incorporated into mixed ubiquitin/NEDD8 chains (Fig 3D). Likewise, mass spectrometry (MS) protein quantification of HA-NEDD8 pulldowns showed enhanced accumulation of catalytically inactive USP8 as compared to HA-NEDD8 pulldowns performed with cells expressing WT-USP8-FLAG (Fig. 3E). Of note, no ubl-modifications of USP8 itself were present among K-ε-GG modified peptides (deposited in PRIDE database, identifier PXD024755). Strikingly, UBA52 (ubiquitin), UBE1 (UBA1), and HUWE1, which represent the characteristic E1 and E3 ligase in proteotoxic stress-induced non-canonical NEDDylation, respectively[43–46,48], were significantly enriched in the HA-NEDD8 pulldowns performed with cells expressing USP8(C786A)-FLAG. Quantification of K-ε-GG modified peptides uncovered the enrichment of K11- and K48-linked ubiquitin/ubl-modifications of NEDD8 and ubiquitin in the same pulldown sample, suggesting that chains containing NEDD8 and/or ubiquitin are formed primarily via K11 and K48 linkages (Fig. 3E). Indeed, when we converted the HA-NEDD8 residues K11 and K48 to arginine (R), less HA-NEDD8 was linked to substrates and a significant decrease in ubiquitylation was observed (Fig. S4A). The formation of mixed NEDD8/ubiquitin chains is further supported by the absence of K48- and K63-specific ubiquitylation, as determined using linkage-specific antibodies, and by the lack of significant sumoylation (Fig. S4B). Moreover, NEDD8 (K22 and K33) and ubiquitin (K63) modifications were present solely in the HA-NEDD8 pulldowns performed with lysates containing USP8(C786A)-FLAG (deposited in PRIDE database, identifier PXD024755). In agreement with non-canonical NEDDylation, the NAE1 inhibitor MLN4924 inhibited Cullin NEDDylation, but not NEDDylation caused by overexpression of USP8(C786A)-FLAG (Fig. 3F). In line with previous data showing that USP8 does not catalyze deNEDDylation[49], USP8 formed a covalent linkage solely with a ubiquitin-propargylamide (PRG) probe[50,51], but not with NEDD8-PRG, which exclusively targeted the deNEDDylating enzyme SENP8[52] (Fig. 3G). These results indicate that overexpression of catalytically inactive USP8 not only induces proteotoxic stress, but that USP8 also cleaves ubiquitin in arising mixed ubiquitin/NEDD8 chains.

The identification of NEDD8-modified substrates besides cullins has been hampered by the lack of efficient MS approaches that can distinguish between ubiquitylation and NEDDylation. Thus, strategies involving the exchange of C-terminal amino acid residues in NEDD8 to alter cleavage of linkage sites by trypsin or LysC proteases prior to MS analysis have been applied[47,53,54]. To confirm ubiquitin/NEDD8 hybrid chain formation (versus parallel chain formation on the same substrate or even non-covalent interactions during pulldown), and to identify ubiquitylation sites on NEDD8 and vice versa, we co-expressed USP8(C786A)-FLAG (and WT-USP8-FLAG) with HA-tagged WT NEDD8 and a variant of NEDD8, in which the C-terminal LRGG sequence was changed to LTGG. NEDD8(LTGG) is still linked to substrates (Fig. 3H, left panels), but trypsin cannot generate the K-ε-GG remnants on which MS identification of the modification sites is based[54]. Following HA-pull down, MS analysis revealed that both, WT-NEDD8(LRGG) and NEDD8(LTGG) were modified on K48, indicating that NEDD8(K48) represents an ubiquitylation site (Fig. 3H, right panel). Vice versa, modification of ubiquitin at K63 was detected in the presence of exogenously expressed NEDD8(LRGG), but not NEDD8(LTGG), indicating that ubiquitin is NEDDylated at K63. In line with USP8-mediated cleavage of ubiquitin in ubiquitin/NEDD8 mixed chains, modification of substrates was diminished by recombinant USP8 (Fig. 3I). Strikingly, *in vitro* DUB cleavage assays with recombinant USP8 and synthetic K48-ubiquitinated NEDD8 or K48-diubiquitin verified that USP8 cleaves ubiquitin/NEDD8 branches even more efficiently than K48-diUb (Fig. 3J).

These data support the conclusion that the absence of USP8 activity promotes proteotoxic stress, as evidenced by the formation of mixed ubiquitin/NEDD8 chains. Additionally, our biochemical results reveal a direct role of USP8 in cleaving these heterotypic chains.

### USP8 controls proteostasis and survival of primary B cells

In line with the finding that exogenous expression of catalytically inactive USP8 induces proteotoxic stress, OHT-induced USP8 deletion in primary *Usp8*^f/f^*Rosa26*-CreERT2 B cells promoted overall ubiquitylation - likewise indicative of proteotoxic stress - and accelerated apoptosis (Fig. 4A). Concordantly, a mild increase in JNK activation was observed. Likewise, Caspase 3 and Caspase 8 cleavage was detected, accompanied by loss of the full-length proteins (Fig. 4B).

**Fig. 4.**
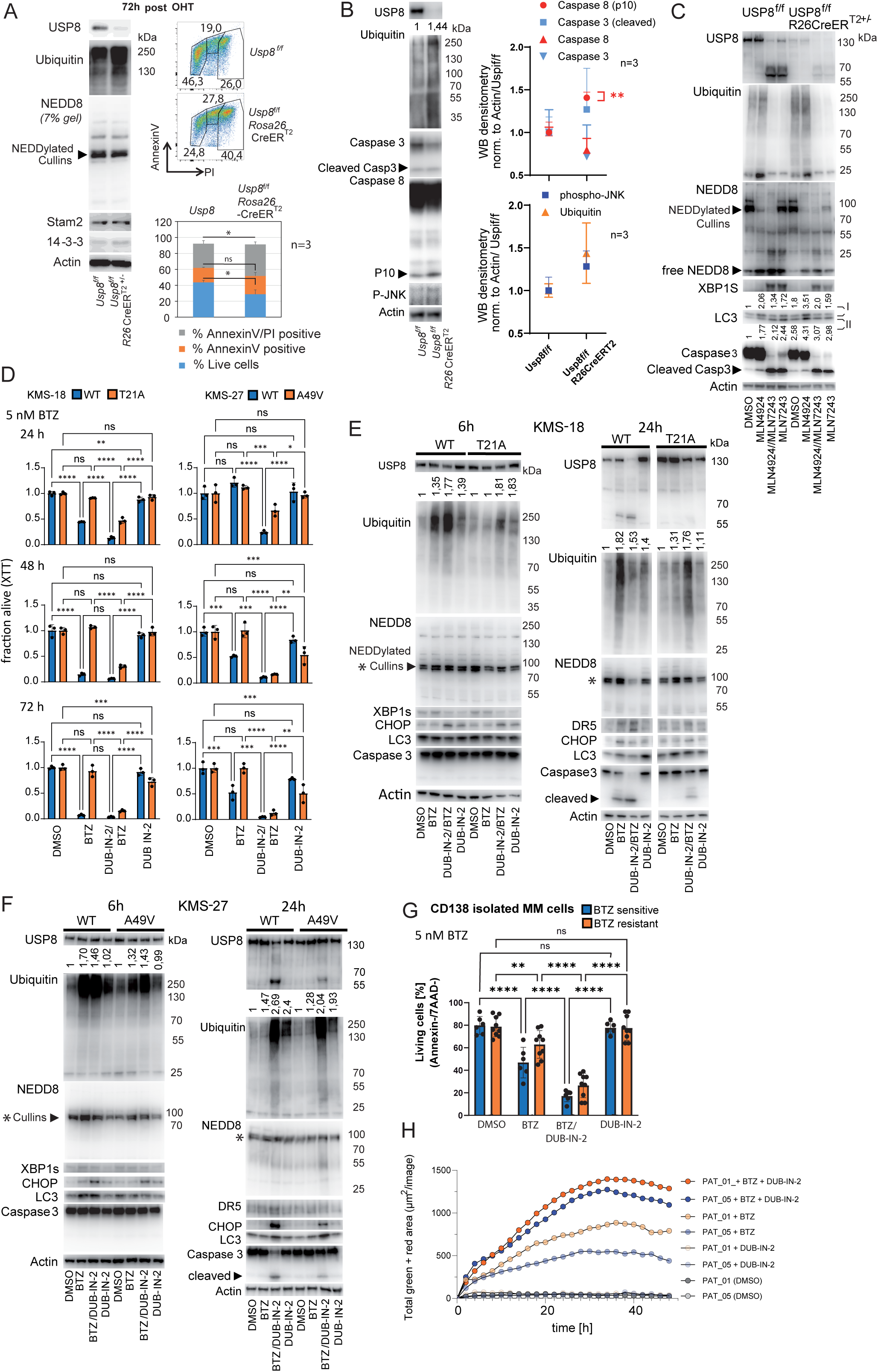
USP8 depletion and BTZ/DUB-IN-2 treatment induce proteotoxic stress and apoptosis in proliferating B and MM cells, respectively. DUB-IN-2 resensitizes resistant MM cells to BTZ. (A) *Usp8*^f/f^ and *Usp8*^f/f^*Rosa26*-CreERT2 splenic B cells were expanded in medium containing IL-4 and CD40 for 24 h followed by OHT addition and further expansion for 72 h. Lysates were subjected to WB analysis (left panels). In parallel, apoptosis was detected by FACS analysis (right panels, n=3, two-tailed unpaired t-test, mean +sd). (B) *Usp8*^f/f^ and *Usp8*^f/f^*Rosa26*-CreERT2 splenic B-cell lysates generated as in (A) were subjected to WB with the indicated antibodies and expression of modified proteins was quantified (n=3, two-tailed unpaired t-test, mean+/-sd). (C) *Usp8*^f/f^ and *Usp8*^f/f^*Rosa26*-CreERT2 splenic B-cells were expanded as in (A) and treated with the inhibitors MLN4924 (NAE inhibitor, 500 nM) and/or MLN7243 (UBA1 inhibitor, 500 nM) as indicated for 16 h before lysis and WB analysis. LC3 I/II quantification is indicated. (D) Cell viability of KMS-18 or KMS-27 WT cells and BTZ-resistant mutant MM cells (KMS-18-T21A/KMS-27-A49V) treated with BTZ (5 nM), DUB-IN-2 (25 µM), or a combination of both for the times indicated (n=3, 2-way ANOVA, Tukey’s post hoc test, mean+/-sd). (E, F) WB analysis of KMS-18/KMS-18-T21A (E) or KMS-27/KMS-27-A49V (F) MM cells after exposure to BTZ (5 nM) and/or DUB-IN-2 (25 µM) for the times indicated (top). (G) BM cells from MM patients defined as “BTZ resistant” (n=9) or “non-resistant” (n=6) were incubated with solvent (DMSO), BTZ (5nM), DUB-IN-2 (25µM), or BTZ+DUB-IN-2 for 24 hours and viability was measured by AnnexinV/7AAD staining. (2-way ANOVA, Tukey’s post hoc test, mean+/-sd; two sided t-test was used to compare the response to BTZ alone across groups, p=0.03). (H) MM cells from selected patients were stained with CytoLight Rapid Green and treated with inhibitors as in (G). Ongoing staining of apoptotic cells was performed with Annexin V Red Dye. WB results in Fig. 4 are representative of at least 2 independent experiments. Quantification of ubiquitin modification shown on top of each blot represents the average of at least 2 independent experiments normalized to actin and DMSO controls. Source data are provided as source data file.

Remarkably, hyperNEDDylation (indicative of mixed ubiquitin/NEDD8 chains) was not detected upon USP8 deletion in these primary B cells. As induction of hyperNEDDylation may require sufficient amounts of free NEDD8[44], we treated USP8-depleted primary B cells with MLN4924 to inhibit the E1 for canonical Cullin NEDDylation. This treatment increased free NEDD8 levels, but also did not cause significant hyperNEDDylation (Fig. 4C). These data suggest that mixed chains are formed only in specific cell-types. Alternatively, enzymes other than USP8 may remove mixed ubiquitin/NEDD8 chains from substrates.

In primary B cells, USP8 depletion leads to induction of autophagy, shown by an increase in the lipidated form of LC3 (II), and inhibition of autophagic flux, indicated by accumulation of LC3 (Fig. 4C).

To evaluate the role of USP8 in ER stress, XBP1s expression was monitored upon USP8 depletion[55]. ER stress was detected if UBA1-mediated ubiquitylation is inhibited by MLN7243, but not upon USP8 depletion (Fig. 4C). Interestingly, UBA1 inhibition triggers USP8 cleavage, correlating with Caspase 3 activation (Fig. 4C).

In summary, these mechanistic studies in B cells following USP8 deletion indicate that inhibition of USP8 function leads to increased protein ubiquitylation and proteotoxic stress-induced apoptosis, but does not affect ER stress.

### DUB-IN-2 controls proteostasis and survival of MM cells in synergy with BTZ

The critical role of USP8 in preventing proteotoxic stress described above and the identification of *Usp8* as a vulnerability gene prompted us to further evaluate its function in MM. This disease is characterized by clonal expansion of malignant PCs in the bone marrow, which are highly susceptible to disturbed protein homeostasis[1,24]. Thus, we speculated that inhibition of USP8 can enhance proteotoxic stress, ultimately resensitizing proteasome inhibition resistant malignant MM cells. We thus initially employed the small molecule DUB-IN-2, which has been reported to specifically inhibit USP8 and was used in multiple studies to evaluate the role of USP8[10–12,18]. As a well-defined MM cell model for BTZ resistance, we took advantage of KMS-18 and KMS-27 MM cells, for which BTZ-resistant variants with well-defined mutations in the BTZ binding pocket of the proteasomal subunit PSMB5 exist (T21A or an A49V substitution, respectively)[56].

We treated KMS-18 and KMS-27 MM cells, as well as their BTZ-resistant variants with BTZ, DUB-IN-2, or a combination of both. While neither BTZ nor DUB-IN-2 sufficiently affected KMS-18-T21A or KMS-27-A49V viability, and DUB-IN-2 did not inhibit viability of WT KMS-18/KMS-27 cells, the combination of both drugs drastically reduced cell viability of both WT and BTZ-resistant cells (Fig. 4D). To overcome BTZ resistance of KMS-18-T21A cells, a DUB-IN-2 concentration of 25 µM was required (Fig. S5A). We observed a similar synergistic inhibitory effect with a combination of the proteasome inhibitor CFZ and DUB-IN-2 in both KMS-27 WT and KMS-27-A49V cells, which also exhibit CFZ resistance[56] (Fig. S5B). Given the limited single-agent efficacy of DUB-IN-2, synergy scores were determined based on a full dose–response matrix (Tables S2 and S3) and evaluated using SynergyFinder Plus[57], with positive scores indicating synergy (Tables S4 and S5). The BTZ and CFZ combination treatments with DUB-IN-2 also led to excessive accumulation of polyubiquitylated proteins within 6 hours (Fig. 4E/F and Fig. S5C). In contrast, increases in overall NEDDylation were induced only in KMS-27 and KMS-27-A49V cells after 24 hours of BTZ/DUB-IN-2 combination treatment underscoring the cell type-specific regulation of this response (Fig. 4F). Similar to UBA1 inhibition in primary B cells, 24h exposure to BTZ led to USP8 protein degradation and caspase 3 activation in KMS-18 WT cells, with a clear correlation between these effects. These were enhanced by DUB-IN-2 co-treatment in all cell lines (Fig. 4E/F, right panels). Consistent with the known role of proteasome inhibition in triggering apoptosis through ER stress in MM cells[58], similar patterns were seen with the upregulation of the ER stress marker CHOP and its target, death receptor 5 (DR5)[55]. Of note, CHOP was also upregulated after 6 hours of treatment with DUB-IN-2 alone, while co-treatment with BTZ (or CFZ) and DUB-IN-2 was the most effective (Fig. 4E/F, left panels, Fig. S5C). XBP1s expression did not parallel upregulation of CHOP in the presence of DUB-IN-2 and may therefore be cell type-dependent and under tight control in MM cells upon proteotoxic stress to promote pro-apoptotic functions as opposed to pro-adaptive functions[59,60]. Autophagy induction, as indicated by the accumulation of the lipidated form of LC3, was not observed (Fig. 4E/F and Fig. S5C).

To complement these findings from KMS-cells cell lines with those from patient derived MM cells exhibiting less well-defined resistance mechanisms, the BTZ/DUB-IN-2 combination treatment was applied to MM cells isolated from 9 patients who either insufficiently responded to BTZ-containing induction therapy or lost their response during BTZ-containing treatment (defined as “BTZ-resistant”) and 6 patients responding well to BTZ-containing induction therapy (defined as “BTZ non-resistant”). The two groups showed a significant difference in the response to BTZ (p=0.03) (Fig. 4G). The combination treatment of BTZ and DUB-IN-2—using BTZ at the concentration that showed the greatest differential effect between ‘BTZ-resistant’ and ‘BTZ-non-resistant’ cells, and DUB-IN-2 at the highest non-toxic concentration (Fig. S5D)—effectively inhibited the viability of these MM cells, further supporting its potential therapeutic relevance (Fig. 4G/H, Table S6).

In summary, our findings suggest that DUB-IN-2 may overcome BTZ and CFZ resistance by enhancing ER stress responses, offering potential therapeutic application.

### DUB-IN-2 impacts MM cell viability independently of USP8

To assess whether consequences of USP8 knockdown in MM cells parallel the effects seen with DUB-IN-2 treatment, we depleted USP8 in KMS cell variants with siRNA. In contrast to the combined BTZ/DUB-IN-2 treatment, combined USP8 depletion/BTZ treatment exhibited an additive or rather mild synergistic effect on cell viability, requiring 3-fold higher BTZ concentrations (Fig. 5A, Bliss scores[61] are listed in Table S7). Notably, KMS-27-WT/A49V cells were more susceptible than KMS-18-WT/T21A cells. In contrast to DUB-IN-2 treatment, USP8 depletion did not induce CHOP or enhance BTZ-induced CHOP expression as an indicator of ER stress, but it induced Caspase 8 and Caspase 3 activation (Fig. 5B). These results suggested that DUB-IN-2 may affect BTZ-resistant MM cell viability independently of USP8 inhibition and raised concerns about its specificity.

**Fig. 5.**
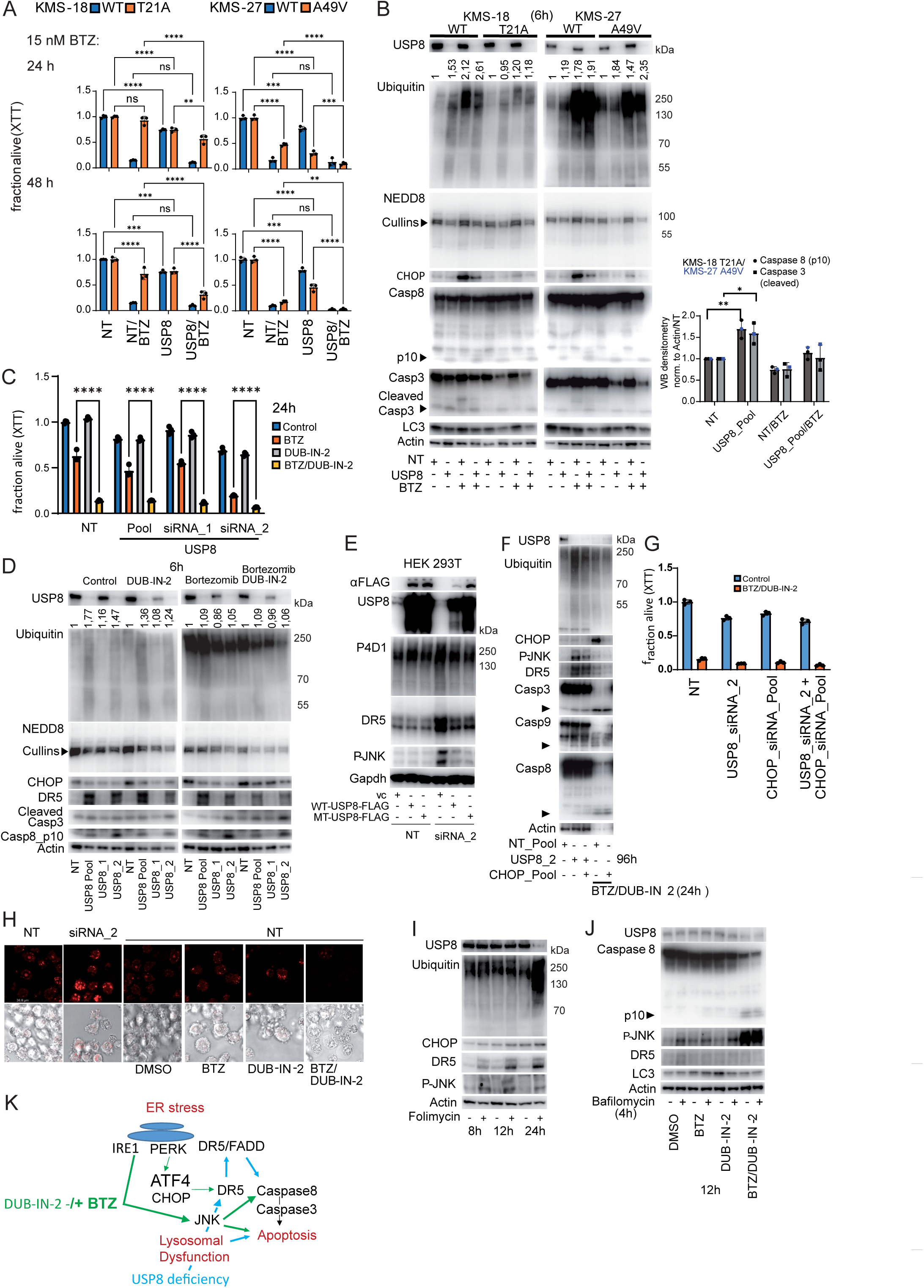
DUB-IN-2 exerts USP8-independent effects on cell viability. (A) Viability of KMS-18/KMS-27 WT and mutant MM cells transfected with pooled (4 siRNAs) non-targeting (NT) or USP8 siRNA for 72 hours followed by BTZ (15 nM) treatment as indicated (n=3, 2-way ANOVA, Tukey’s post hoc test, mean+/-sd). (B) MM cells transfected as in (A) for 96 hours, followed by treatment with BTZ for 6 hours and WB analysis. Caspase processing was quantified (n=3, two-tailed unpaired t-test, mean+/-sd; significant results indicated). (C) Viability of KMS-18-T21A MM cells transfected with pooled NT siRNA, or pooled/individual USP8 siRNA_1/2 for 72 hours followed by 24 h BTZ (25 nM) and/or DUB-IN-2 (25µM) treatment (n=3, 2-way ANOVA, Tukey’s post hoc test, mean+/- sd). (D) KMS-18-T21A MM cells transfected as in (C), treated with BTZ (25 nM) and/or DUB-IN-2 (25µM) for 6 h, analyzed by WB. (E) 293T cells were transfected with USP8-FLAG (WT or silent mutations in the siRNA2 target sequence). USP8 siRNA_2 was transfected after 24h, lysates for WB were generated after 96h. (F) WB analysis of KMS-18-T21A MM cells transfected with pooled NT siRNA, USP8_siRNA_2, pooled CHOP-specific siRNA, or USP8- and CHOP-specific siRNA for 72 h followed by 24 h BTZ/DUB-IN-2 (25 nM/25 µM) treatment as indicated. Arrows indicate cleaved caspases. (G) Viability of KMS-18-T21A cells transfected as in (F) followed by 48 h inhibitor treatment (n=3, mean+/-sd). (H) KMS-18-T21A cells transfected with pooled NT siRNA or USP8 siRNA_2 for 96 h, and treated with BTZ (25 nM) and/or DUB-IN-2 (25µM) for 24 h, as indicated, analyzed by AO staining. Results representative of 5 pictures from 2 independent experiments, respectively. (I) KMS-18-T21A cells treated with Folimycin (100 nM) as indicated, analyzed by WB. (J) WB analysis of KMS-18-T21A MM cells treated with BTZ and/or DUB-IN-2 for 12 h. Bafilomycin (200 nM) added 4 h prior to lysis. (K) Differentially regulated apoptosis pathways triggered by proteotoxic stress upon USP8 depletion (blue) or DUB-IN-2/BTZ treatment (green). Green line thickness indicates DUB-IN-2 effect dependence on BTZ co-treatment. WBs in Fig. 5 are representative of at least 2 independent experiments. Ubiquitin quantification as in Fig. 4. Source data provided.

We therefore evaluated the specificity of DUB-IN-2 in principle. To this end we depleted USP8 in KMS-18-T21A cells using siRNA followed by BTZ, DUB-IN-2, or BTZ/DUB-IN-2 treatment. If DUB-In-2 would target USP8 within this context, we expected a loss of the effect as the target is no longer present (Fig. 5C). USP8 was depleted efficiently with pooled siRNA or siRNA_2, whereas siRNA_1 was less effective (Fig. 5D). Reduced cell viability caused by USP8 depletion and BTZ treatment was further decreased by DUB-IN-2, suggesting that DUB-IN-2 acts independently of USP8. (Fig. 5C).

As these results questioned the specificity of DUB-IN-2 as a USP8 inhibitor, we continued to analyze the consequences of siRNA mediated USP8 depletion. Although USP8 depletion did not increase CHOP expression, its transcriptional target DR5 was strongly upregulated and correlated with caspase 8 activation (Fig. 5D). Partial depletion with siRNA_1 was not sufficient to trigger these effects. DUB-IN-2 and/or BTZ treatment for six hours in the absence of USP8 knockdown had no effect on DR5 expression or caspase activation. However, all treatments enhanced protein ubiquitylation (Fig. 5D). To exclude potential USP8 siRNA off-target effects, we rescued USP8 expression upon USP8 siRNA_2 depletion in 293T cells. Both exogenous WT USP8 and USP8 harboring a silent mutation in the siRNA_2 target sequence rescued USP8 depletion-induced hyper-ubiquitylation, DR5 upregulation, and JNK activation. Of note, JNK activation has previously been shown to act upstream of DR5 transcriptional activation[62] (Fig. 5E) and thus may represent a CHOP-independent pathway of DR5 induction upon USP8 depletion. To further analyze whether DR5 in this situation is indeed upregulated independently of CHOP, we prevented potential CHOP induction in KMS-18-T21A cells by siRNA treatment and analysed DR5 expression after USP8 knockdown. Results further validated that DR5 is upregulated in a CHOP independent manner. (Fig. 5F). Moreover, CHOP was not required for cell death induced by combined BTZ/DUB-IN-2 treatment (Fig. 5F and 5G). In agreement with USP8 depletion-induced JNK activation, which triggers apoptosis via both extrinsic and intrinsic pathways[63], caspase 3, 9, and 8 were activated upon USP8 depletion (Fig. 5 F).

As an ESCRT component, USP8 is involved in lysosomal trafficking[3]. To further investigate the effects of USP8 depletion versus DUB-IN-2 treatment, we analyzed lysosomal integrity in KMS-18-T21A and KMS-27-A49V cells, that have been either depleted of USP8 or treated with BTZ and/or DUB-IN-2. Lysosome enlargement, indicative of impaired lysosomal protein degradation, was observed exclusively following USP8 depletion but not after treatment with DUB-IN-2, BTZ or a combination of both (Fig. 5H and Fig. S5E). We next treated KMS-18-T21A cells with folimycin to chemically induce lysosomal dysfunction. Similar to USP8 depletion, folimycin strongly induced JNK phosphorylation and DR5 expression with little effects on CHOP expression or overall protein ubiquitylation, which was only enhanced after 24 hours (Fig. 5I). These results indicate that USP8 depletion impairs lysosomal degradation, a defect not observed with DUB-IN-2 treatment, thereby raising further doubts about the inhibitor’s specificity for USP8 in this context.

To rule out the possibility that DUB-IN-2 affects autophagosomal protein degradation, we treated KMS-18-T21A cells with bafilomycin in the absence or presence of DUB-IN-2, BTZ or both inhibitors. Bafilomycin inhibits autophagosomal/lysosomal fusion allowing stabilization of the lipidated form of LC3, which was not altered upon inhibitor treatment. Consistent with the pro-apoptotic function of IRE1 in ER stress[59,60], JNK and caspase 8 were strongly induced after 12 h of BTZ/DUB-IN-2 treatment, whereas DR5 expression remained unchanged at this time point (Fig. 5J). Taken together, these results indicate that apoptosis in MM cells treated with BTZ and DUB-IN-2 is mediated by (ER stress-induced) JNK activation in a DR5-independent manner, whereas USP8 depletion triggers JNK activation, DR5 upregulation, and apoptosis predominantly by compromising lysosomal function[64] (Fig. 5K). The engagement of distinct responses upon USP8 silencing, including lysosomal damage and the absence of CHOP upregulation, clearly indicates that DUB-IN-2 exerts USP8-independent effects on apoptosis.

The observed profound differences in cell death inducing mechanisms prompted us to reevaluate USP8 inhibition by DUB-IN-2 and assess potential inhibition of other DUBs. Therefore, a commercially available DUB profiler assay containing more than 40 different recombinant DUBs including USP8 (Ubiquigent) was conducted. DUB-IN-2 was tested at concentrations ranging from 30 nM to 100 µM for each DUB. Surprisingly, no significant inhibition of USP8 or any other DUB was observed, demonstrating that DUB-IN-2 has no substantial inhibitory activity towards USP8 (Fig. 6A/B) or any other DUB tested. Since recombinant USP8 expressed in insect cells was used in the original paper reporting DUB-IN-2 as a USP8 inhibitor [18], we expressed USP8 in SF21 cells and performed a classical deubiquitination assay. To exclude potential contamination from insect cell derived deubiquitinases, catalytically inactive USP8 was used as a control. *In vitro* testing using both, USP8 expressed in insect cells (Fig. 6C/D), and bacterially expressed USP8 (Fig. 6E), did not show DUB-IN-2-mediated inhibition of USP8. Furthermore, the previously reported structurally related USP8 inhibitors DUB-IN-1 and DUB-IN-3[18], and the more recently described inhibitor DC-U4106[19], did not inhibit USP8-mediated deubiquitination (Fig. 6E). In contrast, the dual OTUB1/USP8 inhibitor 61 (OUI 61) affected USP8 activity at concentrations as low as 500 nM (Fig. 6E/F). OUI61 induced significant cell death at 1 µM, but this effect was independent of BTZ treatment and caspase activation, and was not accompanied by the accumulation of ubiquitylated proteins. These results strongly suggest that its toxicity is unrelated to USP8 inhibition (Fig. S6A/B). Finally, DUB-IN-2 had no direct impact on proteasome activity (Fig. S6C). These results unequivocally show that DUB-IN-2 does not inhibit USP8 and corroborate our finding that combined DUB-IN-2/BTZ treatment affects MM viability independently of USP8.

**Fig. 6.**
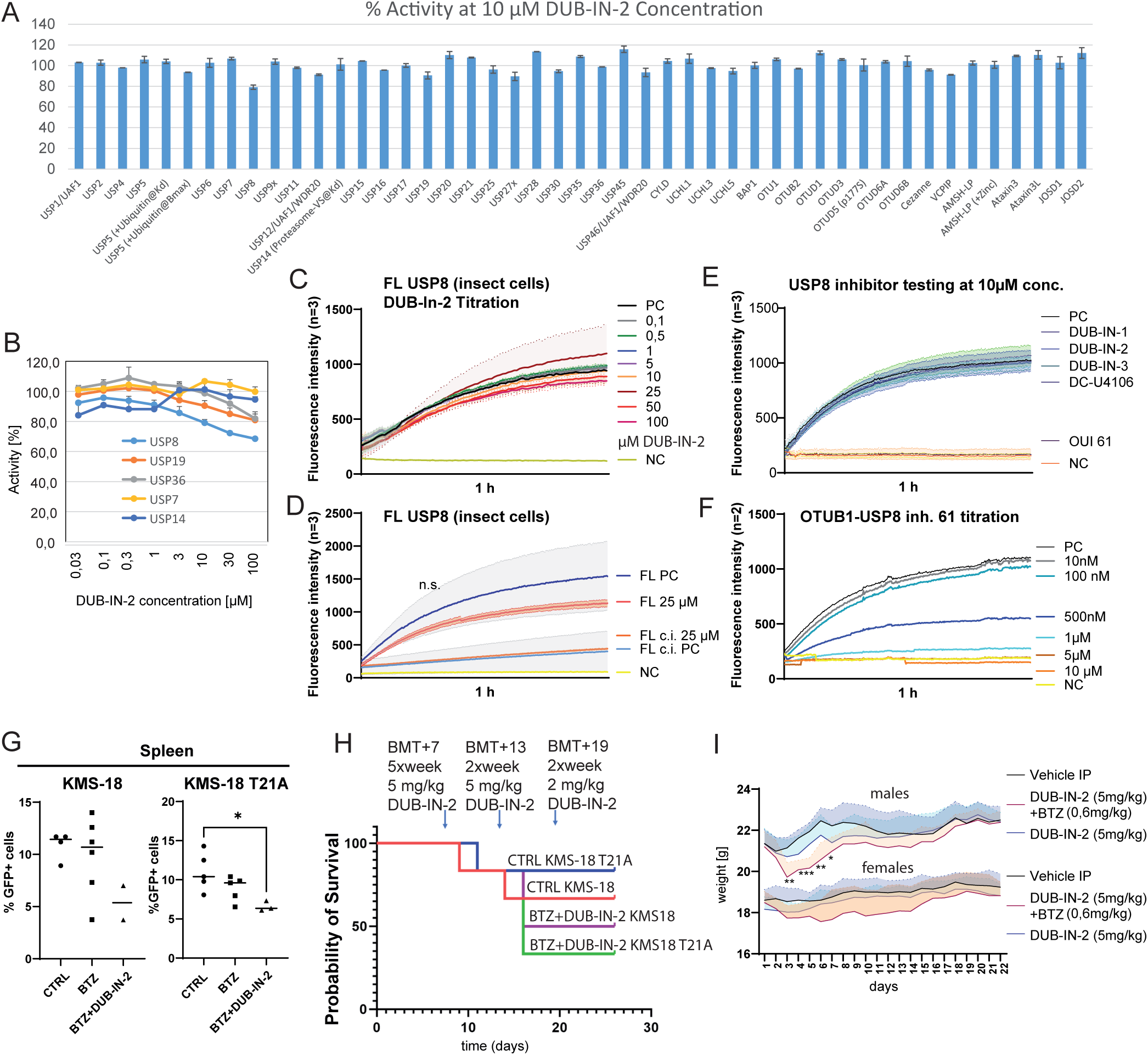
Therapeutic potential of DUB-IN-2 in MM. (A) Evaluation of the selectivity of DUB-IN-2 (10µM) across the DUB*profiler* panel (Ubiquigent). +Ubiquitin@kd/+Ubiquitin@Bmax: Enzyme activated half-maximally/maximally with additional ubiquitin; Proteasome-V5@kd: Enzyme activated with proteasome-V5. (Mean of duplicate experiments +/-sd). (B) Effect of different DUB-IN-2 concentrations on selected DUBs. (Mean of duplicate experiments +/-sd). (C) Evaluation of different DUB-IN-2 concentrations for inhibiting USP8 expressed in insect cells using ubiquitin-rhodamine as a substrate (n=3; PC, positive control; NC, negative control (no enzyme)). (D) Assessment of DUB-IN-2 (25 µM) effects on USP8 and a catalytically inactive (c.i.) variant (C786R) purified from insect cells (n=3). (E) Assessment of commercially available USP8 inhibitors (10 µM) for their effects on bacterially expressed USP8 (n=3). (F) Effect of OTUB1/USP8 inhibitor 61 (OUI61) on bacterially expressed USP8 across different concentrations (n=2). (G) Infiltration of MM cells in spleens of NSG mice treated with either control (0.9NaCl/PEG300 1:1 (v/v)), BTZ (BTZ, 0.6mg/kg, twice a week), or BTZ+DUB-IN-2 (n=6) using 5mg/kg 5 times a week over a period of 13 days, followed by 2 mg/kg twice a week in spleen (1-way ANOVA, Tukey’s post hoc test). (H) Kaplan Meier plot showing the toxicity of the DUB-IN2/BTZ combination treatment (5mg/kg). (I) Weight curves of C57BL/6J mice (n=5) IP injected with 0.9NaCl/PEG300 1:1 (v/v)), BTZ (BTZ, 0.6mg/kg twice a week), or BTZ combined with DUB-IN-2 using 5mg/kg 5 times a week over a period of 22 days. (T-test: combination treatment compared to vehicle control). Source data are provided as source data file.

However, as shown above, DUB-IN-2 was very effective in overcoming BTZ resistance in MM cells. We thus tested the efficacy of BTZ/DUB-IN-2 combination treatment in an *in vivo* transplantation model. To this end, we reconstituted NSG mice with KMS-18 or KMS-18-T21A cells followed by treatment with BTZ alone or a combination of BTZ and DUB-IN-2. Although KMS-18 parental and mutant cell infiltration in the BM was poor within our experimental time frame (Fig. S6D), we detected stable infiltration in the spleen and observed a clear trend of reduced malignant cell expansion upon combined BTZ/DUB-IN-2 treatment in KMS-18 and KMS-18-T21A transplanted mice (Fig. 6G). However, considerable toxicity of the combination treatment was observed (Fig. 6H). In this experimental setup, transplantation was performed upon irradiation of severely immune-compromised NSG mice. To evaluate toxicity of DUB-IN-2/BTZ combinatory treatment in a less vulnerable setting, compounds were applied to non-irradiated, immune-competent mice. Notably, even under more stringent dosing conditions, combined BTZ/DUB-IN-2 treatment did not impact the viability of these mice, and only a transient drop in body weight was observed (Fig. 6I). Moreover, no significant gross pathological alterations were reported upon treatment (Bienta Study Report T011125a). These results diminish concerns about general toxiticity.

In summary, we thoroughly characterized the consequences of USP8 depletion within the B cell compartment of the whole organism and defined its fundamental role in proteostasis. We show that DUB-IN-2 does not represent a specific USP8 inhibitor, but has significant anti MM activity in synergy with BTZ. Our findings strongly suggest that both the specific inactivation of USP8 and treatment with DUB-IN-2 can be exploited therapeutically to overcome BTZ resistance in MM and trigger apoptotic pathways in B-cell malignancies.

## Discussion

Our data demonstrate that USP8 is critical for development and survival of B cells at early and advanced stages. If *Usp8* is deleted at later stages in *Usp8*^f/f^*Cd19*-Cre mice, allowing B-cell development to progress beyond the pre-B-cell stage, immature, innate-like B cells, GC-like B cells, and PCs with diminished Roquin expression predominate, leading to normal (IgG_1_) or enhanced (IgM and IgG_3_) Ig production upon immunization. Roquin proteins, RNA-binding proteins with N-terminal E3 ligase activity, shape T and B cell-dependent immune responses[31,32] by destabilizing target mRNAs (e.g. *Ifng*[31] and *Icos*[65]) and controling mTor activity[66,67]. Roquin ablation in B cells expands the B-cell population and promotes spontaneous GC activation[68], as seen in *Roquin^san/san^* (sanroque) mice, which develop spontaneous GCs and autoimmunity[69]. Preventing GC formation in sanroque mice did not abolish autoimmune disease, but suggested that the expanded B1 B-cell population and extrafollicular PCs contributed to IgM autoantibody generation[70]. Thus, loss of Roquin may contribute to the heightened activation potential of USP8-deficient splenic B cells, including increased JNK activation. In addition, gene expression data suggest that perturbations of Rho-GTPase signaling may be linked to increased Ca^2+^ signaling upon *in vivo* depletion of USP8. However, the precise mechanisms underlying these effects are hard to define and may, in part, be influenced by altered B-cell composition. They are likely dependent on an intact *in vivo* microenvironment, potentially reflecting the broad biological consequences of USP8 loss – including impaired cell viability.

Initial indication of proteotoxic-stress-induced B-cell defects in the absence of USP8 function originated from our experiments performed with K46Lµm B cells expressing catalytically inactive USP8(C748A). In this setting, endogenous USP8 prevents cell death and a USP8(C748A)–mediated “trapping” mechanism leads to enrichment of substrates of USP8 that are modified by mixed ubiquitin/NEDD8 chains[41]. Hybrid ubiquitin/NEDD8 chains promote both formation of cytosolic aggresome-like bodies coupled to the aggresome-autophagy flux[71] and nuclear protein aggregation of mainly ribosomal proteins to protect the nuclear UPS from dysfunction upon proteotoxic stress[48]. Accordingly, NEDD8 has been identified in multiple disease-related inclusion bodies containing ubiquitin, including Lewy bodies in Parkinson’s disease and neurofibrillary tangles from Alzheimer’s disease patients[72,73] corroborating its potential role in antagonizing proteotoxic stress. In accordance with other studies that investigated classical and/or proteotoxic stress-induced substrate NEDDylation[46,47,53], ubiquitin(K63) NEDDylation emerges as a novel regulatory signature specific for proteotoxic stress and functional USP8 deficiency. Moreover, for the first time we identified K48 ubiquitylation of NEDD8 as a USP8 protease target.

Our data indicate that deregulated proteostasis is the main cause of cell death in proliferating USP8-deficient B cells leading to the observed developmental defects. Using MM as a cancer model with high susceptibility to proteasome inhibition and high rates of relapse, we show that the promotion of proteotoxic stress by the compound DUB-IN-2 or USP8 knockdown can be exploited for translational applications. Lack of accumulation of mixed NEDD8/ubiquitin chains in most cell types upon depletion of USP8 might originate from cell type-specific mechanisms. This aligns with the sensitivity of primary B cells and MM cells to proteotoxic stress caused by the absence of USP8, with both cell types lacking UPS-protective mixed chains[48].

In particular, BTZ resistant MM cells could be efficiently targeted with a combination of DUB-IN-2 and BTZ, where DUB-IN-2 alone had little effect on cell viability. In that respect, a caveat of our MM patient cohort is, that most patients defined as BTZ-resistant received a BTZ-containing triple induction therapy. Therefore we cannot rule out that some of those patients actual responded to BTZ but were resistant to one of the two other substances. Samples from patients receiving only BTZ and dexamethasone as MM treatment were rare at our department, which is why we chose to use the current approach. Moreover, in most current treatment regimens BTZ/dexamethasone are used as combination partners for other, modern drugs like CD38-or BCMA-antibodies as well as with selinexor or panobinostat.

MM cells strongly depend on unfolded protein response (UPR) signaling to manage the constant overproduction of proteins, which is essential for their survival and proliferation[60]. Our findings indicate that both BTZ[58] and DUB-IN-2 elicit the apoptotic ER stress response synergistically by targeting the UPS as the “Achilles heel” of MM, via a pathway that is independent of USP8. In contrast, USP8 depletion culminates in lysosomal dysfunction and affects MM cell viability independently of BTZ treatment. Complementary i*n vitro* assays with recombinant USP8 (expressed in bacteria or insect cells) provide additional evidence that DUB-IN-2 does not target USP8 activity. Moreover, DUBprofiler screening and proteasome activity testing did not identify any other DUBs or the proteasome as direct targets of DUB-IN-2. Based on these observations, previous studies where DUB-IN-2 was employed as a USP8 inhibitor would need to be carefully reinterpreted.

However, *in vivo* testing of combined BTZ/DUB-IN-2 treatment in a mouse transplant model as well as on patient samples with undefined resistance mechanisms encourages further studies into exploitation of DUB-IN-2 or derivatives for combination therapies against MM. With regard to translational approaches, it is important to consider that MM can induce immune dysregulation, which may impact vulnerability to BTZ/DUB-IN-2 treatment[74]. While testing a genetically engineered MM mouse model would ultimately provide the most definitive *in vivo* validation, the use of such models is limited by the fact that most inhibitors are specifically designed to target human proteins. In the future, identification of the specific target of DUB-IN-2 will be a major task and might facilitate engineering of an effective MM drug.

In addition, our data encourage further investigations into identification of specific USP8 inhibitors to compromise MM cell proteostasis. Likewise, the critical role and prognostic correlation of USP8 activity in various human cancers[10–13] and neurodegeneration[75,76] opens up new avenues to exploit USP8 as a drug target. Yet, because USP8 strongly affects the tumor microenvironment via regulation of PD-L1 stability, TGF-β signaling, inflammatory responses and, CTLA-4 stability[10–12,14], future investigations should involve models that more closely recapitulate the complexity of human cancer. Correlation of the survival rate of patients from the MMRF-COMMPASS study (https://portal.gdc.cancer.gov/projects/MMRF-COMMPASS) with Usp8 expression at time of diagnosis revealed that the survival rate of MM patients with low USP8 expression tended to have improved survival (Fig. S7A). Moreover, USP8 expression was significantly higher in disease stage III compared with stage II (Fig. S7B) However, it needs to be considered that in these correlative data, variations of USP8 RNA rather than protein levels are monitored.

Early embryonic lethality [33] and fundamental problems in T and B cell development ([20] and this study) in conditional and constitutive USP8 knockout mouse models raised considerable safety concerns about complete USP8 depletion. However, classical genetic deletion models only partially resemble pharmacological approaches. Inhibition by small molecules often exhibits bias toward specific cell types and/or depends on the administration route. As the hematopoietic system is readily accessible for pharmacologic applications, diminishing USP8 activity within this compartment should represent a promising approach for anti-oncogenic and immunomodulatory therapeutics.

## Acknowledgements

We’d like to thank S. Feller and S. Adoro for antibodies, Daniel J. Fernandez for technical advice, and L. Riechert, M. Leffler, M. Ditter and T. Bass for technical assistance. F.I. was supported by an Odysseus type 2 grant from the Research Foundation Flanders (FWO) (G0F8616N)), Concerted Research Action grant BOF21/GOA/033 and Starting Grant BOF/STA/202209/011 from Ghent University, and the European Research Council (ERC Consolidator Grant #101089193). F.T. was supported by a fellowship from the Research Foundation-Flanders (FWO) (12AN524N). G.M. was supported by the Marie Skłodowska-CurieEuropean Individual Fellowship (MSCA-IF-2020, ID: 101025176). N.C. was supported by the IMMediate Advanced Clinician Scientist-Program, Department of Medicine II, University of Freiburg Medical Center, funded by the Bundesministerium für Bildung und Forschung (BMBF, Federal Ministry of Education and Research). O.Gr. received funding through GRK 2606 (Project ID 423813989), and under Germany’s Excellence Strategy, through CIBSS - EXC-2189 (Project ID 390939984). M.S. was supported in the framework of the DFG-funded Research Training Group “Tumor-targeted Drug Delivery” grant 331065168. W.W.S. is supported by the DFG under Germany’s Excellence Strategy - EXC-2189 - Project ID: 390939984 and under the Excellence Initiative of the German Federal and State Governments - EXC-294, by a research grant (Project-ID 403222702 - SFB 1381), and in part by the Ministry for Science, Research and Arts of the State of Baden-Württemberg. Lighthouse Core Facility is funded in part by the Medical Faculty, University of Freiburg (Project Numbers 2021/A2-Fol; 2021/B3-Fol) and the DFG (Project Number 450392965). This work was supported by the Deutsche Forschungsgemeinschaft (DFG) grants 423813989/GRK2606, KN590/4-2 and under Germany’s Excellence Strategy (CIBSS–EXC-2189–Project ID 390939984) to K.P.K.

## Author contributions

Conceptualization, A.D. and K.P.K.; Methodology, F.I., M.F., W.W.S., H.O., P.P.G., H.B., and M.S.; Investigation, A.D., F.T., O.G., N.C., M.F., M.C.A., H.B., J.L., and G.J.H.N.; Formal Analysis, A.D., G.M., F.T., and F.I., M.S.; Writing – Original Draft, A.D. and K.P.K.; Funding Acquisition, K.P.K.; Resources, M.P., H.O., P.P.G., B.J., W.W.S., O.Gr., H.B., K.A., M.S.D., and V.H.; Supervision, A.D., F.I., M.S., and K.P.K..

## Methods

### Mice

*Usp8*^f/f^ [33], *Cd19*-Cre (B6.129P2(C)-*Cd19^tm1(cre)Cgn^*/J)[77], *mb1*-Cre (B6.C(Cg)-*Cd79a^tm1(cre)Reth^*/EhobJ)[27], *Rosa26*-CreERT2 (Gt(ROSA)26Sor^tm2(cre/ERT2)Brn^)[78,79], *Rc3h1*^fl/fl^ [68] and *Rc3h1*^fl/fl^ [80] transgenic, and NSG (NOD.Cg-*Prkdc^scid^ Il2rg^tm1Wjl^*/SzJ)[81] mice have been described. Mice used for experiments were 8-12 weeks old and on the C57BL/6 background. *Rosa26*-Cre-ERt2 mice[79] in combination with/without *Rc3h1/2*^fl/fl^ alleles which are transgenes for Roquin-1 [68] and Roquin-2 [80], respectively, were fed by oral gavage with 5 mg tamoxifen (Sigma) in corn oil per dose on two consecutive days with two doses of tamoxifen each day (20 mg total tamoxifen dose per mouse) and mice were killed 3 d after the last gavage. For histology, B-cell enrichment, and if paired T-tests were performed, sex-matched littermates were used. Animals were housed in a specific pathogen-free barrier facility according to the protocols of the Center for Experimental Models and Transgenic Service at the University Clinic in Freiburg, the LMU Munich, or at the RWTH University Aachen. All experiments were approved by national authorities and the ethics review board for animal studies at the University of Freiburg, the LMU Munich, or RWTH University Aachen.

### Repeated dose toxicity study of DUB-IN-2 and BTZ

Evaluation of the general toxicity of two compounds, DUB-IN-2 (5 mg/kg, 5x per week IP) alone, and in combination with BTZ (0.6 mg/kg, 2x per week IP) was carried out by BIENTA, Enamine Biology Services, Ukraine. 11-12 weeks old male and female C57BL/6J mice were divided into three groups per sex consisting of five C57BL/6J mice each. The mice were repeatedly IP dosed with vehicle (PEG300 – saline, 50%:50%, v/v), test compound DUB-IN-2 (5х per week) in vehicle or combination of the test compounds DUB-IN-2 (5х per week) and BTZ (2x per week) for 22 consecutive days. Animals were weighed on the day of randomization and every day during the study (from 9 am to 10 am). Observations for mortality were performed once a day during the dosing period. The toxic effects of compounds were evaluated during the study (22 days) by visual observations according to the standard procedures and FELASA recommendations. Autopsy with gross pathology examination was performed on Day 22 of the study. All experimental animal procedures performed were approved by the local Institutional Animal Care and Use Committee of BIENTA LTD (BACUC, Approval number: EN-TOX-AP-211124) and in adherence with the European Convention for the Protection of Vertebrate Animals used for Experimental and other Scientific Purposes.

### Southern blot analysis

LoxP-flanked and deleted *Usp8* alleles were identified by Southern blot analysis of NcoI fragments of genomic DNA as described[33].

### Cell preparation and flow cytometry

Spleens, lymph nodes, and thymi were mashed into single-cell suspensions in FACS buffer (PBS containing 1-2% FCS). Bone marrow was obtained from tibias and femurs by flushing with FACS buffer. Peritoneal cavity cells were harvested post-euthanasia by i.p. injection of FACS buffer and peritoneal massage. Splenocytes and BM cells were depleted of red blood cells with RBC lysis buffer (Sigma or BioLegend). Cell suspensions were incubated with mouse Fc Block (BD or BioLegend). B cells were isolated from Spleen using biotinylated anti-CD19 antibody (BD Pharmingen, clone ID3) or the BD IMag^TM^ Biotinylated Mouse B Lymphocyte Enrichment Cocktail and BD IMAG^TM^ Streptavidin Particles Plus. T cells were isolated using the BD IMag^TM^ Biotinylated Mouse CD4 T lymphocyte enrichment Cocktail and BD IMAG^TM^ Streptavidin Particles Plus. DAPI or Zombie UV fixable viability dye (Biolegend) were used for live/dead discrimination. Expression of surface and intracellular markers was analyzed with a flow cytometer (Becton Dickinson FACS Calibur, BD FACSCanto II, or BD Fortessa SORP) and FlowJo software. Annexin V/PI staining was performed using the eBioscience Annexin V apoptosis detection kit APC.

### Cell culture, proliferation, activation, and viability

For the proliferation experiments, purified B cells (5×10^4^ per well) were stimulated by incubation with 5 µg ml^−1^ anti-IgM F(ab’) fragment antibody (115-006-020; Jackson Immuno Research) with or without 5 µg ml^−1^ anti-CD40 antibody (553787, BD Pharmingen), with 100 ng/ml MPL-A (Invivogen) or 100 nM CpG (ODN 1668, Invivogen). B-cell proliferation was quantified after 36 h using the Cell Proliferation ELISA, BrdU (colorimetric) from Roche. For short-term stimulation, B cells were starved for 2 h in RPMI 1640 medium containing 0.2% bovine serum albumin without FCS and activated by preincubation with 10 µg ml^−1^ anti-IgM for 30 min on ice and stimulation for the times indicated at 37°C. T cells were starved for 2 h in RPMI 1640 medium without FCS. T cells were activated by pre-incubation on ice for 30 min with soluble anti-mouse CD3ε antibody (10µg ml^−1^; 145-2C11; eBioscience) and anti-mouse CD28 antibody (2µg ml^−1^; 37.51, eBioscience), followed by stimulation for the appropriate time at 37°C with pre-warmed anti-hamster IgG1 antibody (5µg ml^−1^; G94-56, BD Pharmingen). *Ex vivo* expansion of B cells was performed using recombinant murine IL-4 (100 u ml^−1^, Peprotech) and anti-mouse CD40 antibody (5 µg ml^−1^; HM40-3, Biolegend), or anti-IgM F(ab’) fragment (5 µg ml^−1^) and anti-CD40 antibody (2.5 µg ml^−1^). OHT (1 µM, Sigma) was added 24 h post B-cell isolation and stimulation. DUB-IN-2 (25 µM, MedChemExpress), BTZ (5 to 25 nM, Selleckchem), CFZ (3 nM, MedChemExpress), and OTUB1/USP8 inhibitor 61 (Aobious) were added for the times indicated before analysis. Cell viability was analyzed using the XTT cell proliferation kit II (Roche Diagnostics). Absorbance was measured as indicated and normalized to the respective controls.

### Effect of inhibitors on MM cells of patients

BM cells from MM patients were prepared from the pelvic crest. CD138+ MM cells were isolated from the BM using CD138 microbeads (Miltenyi Biotech, Bergisch Gladbach, Germany). The MM cell purity was >90%. Isolated MM cells were co-incubated with solvent (DMSO), Bortezomib 5nM;. DUB-IN-2 25µM or Bortezomib + DUB-IN-2 for 24 hours. DMSO concentrations were adjusted to match across all samples. Negative control incubations contained 0.1% (v/v) DMSO in the respective incubation buffer. Viability of MM cells was measured by AnnexinV and 7AAD staining (BD) according to the manufacturer’s instructions (FACS Canto II, BD).

In some experiments, isolated MM cells were stained with CytoLight Rapid Green (Sartorius, Göttingen, Germany) and seeded in a 96 well plate. Cells were treated with the inhibitors described above and the culture medium was supplemented with Annexin V Red Dye (Sartorius, Göttingen, Germany) to facilitate ongoing staining of apoptotic cells. Green and red fluorescence channels were recorded once every 120 min over a period of 48 hours. The aggregated area of red cells (µm^2^/image) as a measure of cell death was automatically quantified by Incucyte Base Software (Sartorius, Göttingen, Germany).

BM cells were collected and used from left-over material of myeloma patients upon diagnostic bone marrow aspiration, which was approved by the local ethics committee of the University of Erlangen (219_14B, 19-56D).

### Single cell RNA-seq analysis

Single cell RNA-seq analysis was performed with splenic B cells enriched from *Usp8*^f/f^ and *Usp8*^f/f^*Cd19*-Cre mice using the BD IMag^TM^ Biotinylated Mouse B Lymphocyte Enrichment Cocktail and BD IMAG^TM^ Streptavidin Particles Plus (n=3). For single cell library preparation, cells were loaded onto a Chromium Single Cell 3′ G Chip (10X Genomics) according to the manufacturer’s instructions without modifications using Chromium Next GEM Single Cell 3’ Reagents Kit v3.1 (10x Genomics). The 6 samples were multiplexed using Cell Multiplex Oligos (CMOs) for CellPlex reagents. Dual Index Plate NN Set A (10x Genomics) was used for Cell Multiplexing library construction, Dual Index Plate TT Set A was used for Gene Expression library construction according to the manufacturer’s instructions. The cDNA content and size of post-sample index PCR samples was analyzed using a 2100 BioAnalyzer (Agilent). Library quantification was done using NEB Next® Library Quant Kit for Illumina® (New England Biolabs) following manufacturer’s instructions. Sequencing libraries were loaded onto an Illumina NextSeq 1000 P2 flow cell, with sequencing settings according to the recommendations of 10x Genomics (read 1: 28 cycles, read 2: 90 cycles, i7 index: 10 cycles, i5 index: 10 cycles). Cell Ranger v6.1.2 software was used to align the sequencing data to the mouse genome mm10. The indexed mouse genome used for alignment was downloaded from 10x genomics (refdata-gex-mm10-2020-A).

The single cell data were loaded into R and in a Seurat object (v.4) for subsequent bioinformatics analysis. Cells with a high mitochondrial content (> 20%) or low number of genes detected (<200) were discarded. SoupX[82] was used to estimate and remove cells contaminated from ambient mRNA and ALRA[83] was used for data imputation.

The data were normalized with the “LogNormalized” method and scaled. The top 10.000 variable genes were used for the Principal Component Analysis and the UMAP was generated using the top 20 principal components. The Louvain algorithm with resolution=0.8 from the FindClusters function was used to assign clusters to cells in an unbiased way. Cell type annotation was performed using the markers reported by Nguyen et al.[28] and clusters were merged when no relevant difference in marker expression was detected. Other bioinformatics analyses and data visualization were performed with the other Seurat or ggplot2 functions. The data are available at https://www.ncbi.nlm.nih.gov/geo/query/acc.cgi?acc=GSE244334 (access token for reviewers: **sxsdsecavdyxbqj**). Pathway enrichment analysis was performed with ReactomePA v1.46.0 ( <u>10.1039/c5mb00663e</u>). Genes with adjusted p value < 0.1 and log2FC > 0 or <0 were selected to perform the enrichment analysis for pathways upregulated in the *Usp8*^f/f^*Cd19* B cell/PC and control samples, respectively. Pathways that had 70% of overlapping genes were removed from the final results in order to remove redundancy.

### Cell lines, expression constructs, transfection and inhibitor treatment

NP-specific mIgM-BCR–expressing transfectant K46Lμm cells of the Balb C mouse B-cell lymphoma line K46[38,39] were obtained from W. Schamel, Freiburg. The human MM cell lines KMS-18 and KMS-27 and their mutant T21A and A49V derivatives expressing PSMB5 with a substitution in the BTZ binding cleft were described previously[56]. Human and mouse USP8 tagged with FLAG at the carboxyl terminus were expressed via the vector pFLAG-CMV5a and the retroviral expression vector MIEG3, respectively. Retroviral particles were produced in Platinum-E retroviral packaging cells according to standard protocols. Mammalian expression vectors for HA-NEDD8 and HA-SENP8 were purchased from addgene. Point mutations were generated by site-directed mutagenesis with the QuikChange mutagenesis system (Stratagene). HEK293T cells were transfected with FuGENE HD transfection reagent (Promega) and were harvested 36 h post transfection. The NEDD8-activating enzyme (NAE) inhibitor MLN4924 (LKT Laboratories), the UBA1 inhibitor MLN7243 (MedChemExpress), Folimycin (MedChemExpress), BTZ (Selleckchem), and DUB-IN-2 (MedChemExpress) were applied to cells as described. ON-TARGETplus SMARTpool siRNA directed against human USP8 and CHOP, individual siRNAs directed against USP8 (Target Sequence siRNA_1: GGCAAGCCAUUUAAGAUUA; siRNA_2: CCACUAGCAUCCACAAGUA), and ON-TARGETplus Non-targeting (NT) Pool siRNA were purchased from Dharmacon. SiRNA transfections were perfomed using Lipofectamine® RNAiMAX reagent according to manufacturer’s instructions. For rescue of USP8 depletion by siRNA_2, the respective target sequence was mutated as follows: CC<u>C</u>CUAGC<u>U</u>UCCACAAGUA.

### Immunoblot analysis and immunoprecipitation

Cells were lysed in DDM lysis buffer (50 mM Tris-HCl, pH 7.4, 150 mM NaCl, 1 mM EDTA, 50 mM NaF, 1mM sodium vanadate, 1x Roche Complete Protease Inhibitor Cocktail, and 0.2% n-dodecyl-β-D-maltoside), or directly in 4x Laemmli buffer supplemented with 6 M Urea. Immunoprecipitation was performed in DDM lysis buffer with anti-FLAG M2 Affinity Gel (Sigma) or rat monoclonal antibody to hemagglutinin (HA) affinity matrix (3F10; Roche). Immunoblot analysis was performed with antibodies listed in Supplementary Table S8. Quantification of WB band intensities was performed by measuring the integrated density using Adobe Photoshop software.

### Conjugation and deconjugation assays

For the PRG probe conjugation assay, protein lysate was diluted 1:8 in 25 µl assay buffer (50 mM Na_2_HPO_4_, pH7.9, 500 mM NaCl, 10mM DTT) containing 2 µg NEDD8-PRG or 5µM Bt-Ub-PRG and incubated for 2 h at 37°C. To test the deconjugation capacity of USP8 towards mixed ubiquitin/NEDD8 chains, 5µl protein lysate generated with DDM lysis buffer w/o proteinase inhibitors was incubated with 365 ng recombinant USP8 in PBS containing 5mM MgCl_2_ and 2mM DTT for 1 h at 37°C.

### DUB activity analysis, inhibitor testing, and production of recombinant USP8

The selectivity of DUB-IN-2 was evaluated in the DUB*profiler* assay (Ubiquigent) against 48 individual deubiquitinase enzymes, at concentrations of 0.03, 0.1, 0.3, 1, 3, 10, 30, and 100 μM. Values in the presence of compound were compared with those of DMSO controls to give the percentage of remaining activity. DUB-IN-2 did not show any significant autofluorescent properties in the ubiquitin-rhodamine assay up to 100 μM.

Bacterially eypressed full length USP8 was a gift from Emma Murphy (Alzheimer’s Research UK Oxford Drug Discovery Institute, University of Oxford). WT and catalytically inactive His6-tagged USP8 were expressed in Sf21 cells using the MultiBac system[84]. In brief, a His6-tag followed by the HRV 3C recognition site was amplified from the vector pKL-His-3C-USP18[85] (BamHI-His-fwd: AAAGGGATCCATGGGTCACCATCACCACC; His-EcoRI-rev: AAAGAATTCACCCGGACCCTGGAACA) and ligated into pFL vector[86] yielding vector pFL-His-3C. Human USP8 cDNA was amplified (USP8_SalI_FL_fwd: AAAGTCGACGAATGCCTGCTGTGG; USP8_HindIII_FL_rev: AAAAAGCTTTTATGTGGCTACATCAGTTACT) and ligated into the pFL-His3C vector. Constructs for USP8C786R expression were generated using QuikChange site directed mutagenesis (Agilent; USP8C786R_f: CTTAGGAAATACTCGTTATATGAACTC; USP8C786R_rev: GAGTTCATA TAACGAGTATTTCCTAAG). Constructs were transformed into competent E. coli harboring the MultiBac bacmid[86] via electroporation. Sf21 insect cells were transfected with transposed bacmid DNA using X-tremeGene HP DNA transfection reagent (Roche). Primary virus (V0) was harvested after 2–3 days and used to generate secondary virus (V1) by infecting cells at a density of 0.5 × 10 cells/mL. Infected cells were monitored for fluorescence, harvested after 5–7 days, and protein expression was confirmed by SDS-PAGE and WB analysis. For batch purification of USP8 expressed in insect cells, cell pellets were thawed on ice and resuspended in lysis buffer (20 mM Tris, pH 8.0, 1 M NaCl, 1% Nonidet P-40, 10 mM DTT, and 25 mM imidazole). The suspension was incubated on ice for 10 minutes and subsequently sonicated to facilitate cell lysis. The lysate was centrifuged at 10,000 × g for 10 minutes at 4°C to separate cellular debris and DNA. The cleared lysate was then combined with pre-washed INDIGO-Ni agarose resin (Cube Biotech) and incubated with gentle rotation at 4°C for 3 hours to allow binding of His-tagged proteins. The flow-through was discarded, and the resin was washed sequentially with buffers containing increasing concentrations of imidazole (20 mM Tris, pH 8.0, 1 M NaCl, 10 mM DTT, 40–100 mM imidazole) to remove non-specifically bound proteins. Finally, the His-tagged proteins were eluted by incubating the resin with elution buffer (20 mM Tris, pH 8.0, 300 mM NaCl, 10 mM DTT, and 500 mM imidazole) at room temperature for 10 minutes. DUB activity assays with bacterially expressed human FL USP8 (30 nM) or human USP8 expressed in insect cells (75 nM) were performed in assay buffer (20 mM HEPES pH 7.4, 150 mM KCl, 10 mM NaCl, 5% glycerol, 0.2 mg/ml BSA, 5mM DTT (freshly added)) using ubiquitin-rhodamine-morpholine (100 nM) as a substrate. Fluorescence intensity was measured over a period of 1 hour using a Spark® TECAN Reader with an excitation wavelength of 480 nm and an emission wavelength of 520 nm, controlled by Spark Control software v3.2.

### Proteasome activity testing

Proteasome activity was evaluated using fluorogenic substrates specific for chymotrypsin-like (Suc-LLVY-AMC), trypsin-like (Boc-LRR-AMC), and caspase-like (Z-LLE-AMC) activities. The assay was performed with the Proteasome Activity Fluorometric Assay Kit (UBPBio) following the manufacturer’s instructions.

### ELISA, T cell-dependent and -independent Ig responses, cytokine bead array

Basal immunoglobulin levels in mouse plasma were analyzed by Aushon Biosystems, Billerica, MA. *Usp8^f/f^* and *Usp8^f/f^Cd19*-Cre mice have been i.p. injected with 50 µg TNP-KLH (Biosearch Technologies) precipitated in alum (TD response) or with 25 µg TNP-Ficoll (Biosearch Technologies, TI response) and blood was collected at day 0 and days 7, 14, and 28 post injection. TNP-specific IgM, IgG1 and IgG3 levels in the plasma were quantified using the SBA Clonotyping^TM^ System/AP from SouthernBiotech. Detection of serum cytokine levels was performed using Mouse Enhanced Sensitivity Master buffer Kit (562246, Becton Dickinson) along with Flex Kits for TNF (562336), IL-6 (562236) and IL-1beta (562278) according to manufacturer’s recommendations.

### Ca^2+^ flux

B cells were purified, starved of serum as described above and loaded for 30 min at 37°C with Indo-1-AM (5µg/ml) in RPMI 1640 medium without FCS. Stimulation with anti-IgM (5µg/ml; identified above) was carried out in RPMI-1640 medium with 10% FCS. The emission wavelength ratios of intracellular Ca^2+^-bound and unbound Indo-1-AM were analyzed on a LSRII or a BD Fortessa SORP flow cytometer with FACSDiva software (BD Biosciences).

### Immunohistochemistry and acridine orange (AO) staining

Hematoxilin and eosin staining and immune histological staining of splenic paraffin or cryosections with antibodies was carried out with the appropriate antibodies (dilution 1:500, except for CD3: 1:100; anti-B220: 1:200; Ki67: 1:50; PNA: 1:250) (Supplementary Table S8) as described previously[20,87]. In brief, spleens were removed and fixed in 4 % buffered formalin. Paraffin-embedded 3 μm sections were deparaffinized and were incubated in 3 % H_2_O_2_ in methanol and subsequently in 70 % ethanol. Cryosections were fixed for 5 min in ice-cold acetone, followed by incubation with 10% FCS in PBS. Spleen sections were incubated with the primary antibodies overnight. Secondary antibodies and fluorescence labelled streptavidin (1:500) were added for 2 h at room temperature. Images were taken using a conventional fluorescence microscope (Olympus BX-61). Cells were stained in AO (100 ng/ml, Invitrogen) according to manufacturer’s instructions. Live cell imaging was performed at 37°C in a 5% CO2 humidified atmosphere with a Leica SP8 confocal microscope equipped with a 63×/1.40 and 40×/1.25 oil objective (Leica Microsystems) or a ZEISS Celldiscoverer 7 confocal microscope (Lighthouse Core Facility). Excitation was performed using the 488 nm laser; emission was recorded simultaneously in both the GFP range (for nucleic acids) and the Cy5/Alexa647 range (for protonated AO trapped in organelles). Laser settings were kept constant for each experiment to allow direct comparison of signal intensities between images of the same channel. LAS X software (version 3.7.3.23245) was used for confocal imaging with the Leica TCS SP8 X. ZEN 3.4 (Blue Edition) software was used for confocal imaging with the ZEISS Celldiscoverer 7 confocal microscope.

### Generation of stable GFP expressing cell lines

KMS-18 and KMS-18-T21A cell lines were seeded at 1×10^5^ in 200µl medium (RPMI 1640 supplemented with 10% Fetal Bovine Serum, without antibiotics) and mixed with polybrene (16µg/ml, Sigma-Aldrich). Cells were then placed in a 48-well plate (Thermo Fisher Scientific) together with the virus particles (Firefly luciferase-GFP lentivirus, System Biosciences, stock: 21,9 TU/ml, MOI 10). The plate was centrifuged for 2h at 1200g and 32°C. After centrifugation, cells were expanded for 5 days (37 °C and 5% CO2) in the presence of puromycin (2 µg/ml) and the GFP positive cells were isolated by flow cytometry.

### Transplantation of NSG mice

1×10^6^ MM KMS-18 or KMS-18-T21A cells, combined with 5×10^5^ HS27a human stromal cells were transplanted into 9-week-old NOD/Scid/IL2Rγ-/-(NSG) mice (Charles River) after sublethal irradiation with 240 cGy (2,4Gy) using MultiRad 225 (Cu 0,3mm filter). To prevent infections, cotrimoxazol (100 μg/ml) was added to the drinking water for 2 weeks after transplantation. Bortezomib was administered twice per week (i.p.) at a concentration of 0,6mg/kg and DUB-IN-2 solved in 0.9% NaCl/PEG300 1:1 (v/v) was administered 2-5 times per week (i.p.) at concentrations of 2-5 mg/kg. As control we administered 0.9% NaCl/PEG300 1:1 (v/v).

### Chemical synthesis of K48-ubiquitinated-NEDD8 and K48-diubiquitin

K48-ubiquitinated-NEDD8 and K48-diubiquitin were chemically synthesised as described by El Oualid et al.[88]. In brief, Ubiquitin_1-76_-thioester and K48-γ-thiolysine-Ub or NEDD8 were prepared using automated solid phase peptide synthesis. The StBu protected K48γ-thiolysine-Ub or NEDD8 (5 mg) was dissolved in 50 µL DMSO and added to 600 µL 8 M Gdn.HCl/100 mM phosphate buffer at pH 7.6 supplemented with 50 µL 1 M TCEP and reacted at 37 ^0^C. After LC-MS analysis revealed complete deprotection of the thiolysine, 1.25 equivalent of Ub-thioester dissolved in 50 µL DMSO and 50 uL 1M MPAA were added. The pH was re-adjusted to 7.6 and the reaction was performed for 2 hours at 37 ^0^C. Following RP-HPLC purification the appropriate fractions were lyophilized. The lyophylised diUb(l) (1.5 mg) was desulfurized by dissolving in 25 µL DMSO and subsequent dilution into 600 µL 8 M Gdn.HCl/100 mM phosphate buffer at pH 7.6 and 150 µL 1M TCEP. VA044 (25 mg/mL) and GSH (25 mg/mL) were added, the pH was adjusted to 7.0 and the reaction was agitated for 16 hours at 37 ^0^C. Following RP-HPLC purification the appropriate fractions were lyophilized. The lyophilized diUb(l) was dissolved in 25 µL DMSO and diluted into 750 µL 20 mM TRIS, 150 mM NaCl at pH 7.6 and purified on a S75 10/300 sephadex size exclusion column (GE Healthcare). Appropriate fractions were collected and pooled. Spin-filtration (Amicon 3 kD MWCO) was used to concentrate the sample to 3.4 mg/mL, based on nanodrop analysis (M.W.: 17089 Da, ε_280_: 4470 M^−1^cm^−1)^. The aliquots were snap frozen and stored at −80 ^0^C until further use.

### HRMS-measurements

High resolution mass spectra were recorded on a Waters Acquity H-class UPLC with XEVO-G2 XS Q-TOF mass spectrometer equipped with an electrospray ion source in positive mode (source voltage 3.0 kV, desolvation gas flow 900 L/hr, temperature 250°C) with resolution R = 22000 (mass range m/z = 50 2000) and 200 pg/uL Leu-Enk (m/z = 556.2771) as a lock mass. Data processing was performed using Waters MassLynx MS Software 4.1 (deconvolution with MaxEnt1 function).

#### K48-ubiquitinated-NEDD8

HR-MS analysis for C_757_H_1266_N_208_O_234_S_2_ : Deconvoluted ESI MS^+^ (amu) calcd: 17089.5, found 17089.0, rt: 1.35 min.

#### K48-diubiquitin

HR-MS analysis for C_757_H_1258_N_210_O_235_S : Deconvoluted ESI MS^+^ (amu) calcd: 17093.7, found 17095.0, rt: 1.36 min.

### DUB cleavage assays

USP8 (human, full length, Ubiquigent 64-0053-050) was diluted to a final concentration of 40 nM in buffer containing 20 mM Tris-HCl, 150 mM NaCl, pH 7.6 and 5 mM DTT. Subsequently 5 µM synthetic K48-ubiquitinated-NEDD8 or K48-diubiquitin were added. The samples were incubated at 37°C for 30, 60 and 120 minutes. At the indicated time the reaction mixture was diluted with sample buffer (3x) (containing 75% NUPAGE® LDS sample buffer (4x, Invitrogen), 7.5% β-mercaptoethanol and 17.5% water), and loaded on 12% NUPAGE® Novex® Bis-Tris Mini Gels (Invitrogen) using MES-SDS as running buffer. SeeBlue Pre-stained Standard (Invitrogen, LC5925) was used as molecular weight marker. Gels were stained using InstantBlue™ (Expedeon) and scanned using a GE Healthcare Amersham Imager 600.

### Ubiquitylation proteomics

#### Sample preparation for pull down-MS experiments

In the pull down-MS experiments, after immunoprecipitation beads were washed 2 times with DDM lysis buffer and three times with trypsin buffer containing 20 mM Tris HCl, pH 8.0; 2 mM CaCl_2_. The proteins were on-bead digested with 1 µg trypsin (Promega) for 4 hours at 37 °C under agitation. Then, the beads were precipitated and the supernatants were mixed with another 1 µg of trypsin and further digested overnight at 37°C under agitation. Peptides were desalted on reversed phase C18 OMIX tips (Agilent), dried under vacuum in HPLC inserts and stored at −20 °C until LC-MS/MS analysis.

#### Sample preparation for GlyGly(K) sites identification by MS

200 x 10^6^ K46Lµm cells were homogenized in 10 mL urea lysis buffer containing 9 M urea; 20 mM HEPES, pH 8.0. Samples were sonicated by three pulses of 10 s at an amplitude of 20% and centrifuged for 15 min at 16,000 × g at room temperature to remove insoluble components. The protein concentration in the supernatants of each replicate was measured using a Bradford assay (Biorad) and equal protein amounts, each containing 10 mg total protein, were used for further analysis. Proteins in each sample were reduced by adding 5 mM DTT and incubation for 30 min at 55 °C. Alkylation of the proteins was done by addition of 10 mM chloroacetamide and incubation for 15 min at room temperature in the dark. Next, samples were diluted with 20 mM HEPES, pH 8.0 to an urea concentration of 4 M and proteins were digested with 40 µg endopeptidase LysC (Promega) (1/250, w/w) for 4 h at 37 °C. Samples were further diluted with 20 mM HEPES pH 8.0 to a urea concentration of 2 M and proteins were digested with 50 µg trypsin (Promega) (1/200, w/w) overnight at 37 °C. Immunocapture of GG-modified peptides was then performed using the PTMScan® Ubiquitin Remnant Motif (K-ε-GG) Kit (Cell Signaling Technology) according to the manufacturer’s instructions. Briefly, peptides were purified on Sep-Pak C18 cartridges (Waters), lyophilized for 2 days and re-dissolved in 1.4 ml 1x immunoprecipitation buffer supplied with the kit. At this point, aliquots corresponding to 50 µg of digested protein material were taken for shotgun proteomics analysis. Peptides were incubated with the antibody-bead slurry for 2 h on a rotator at 4 °C and after several wash steps, GG-modified peptides were eluted in 100 µl 0.15% TFA and desalted on reversed phase C18 OMIX tips (Agilent). Purified GlyGly-K modified peptides were dried under vacuum in HPLC inserts and stored at −20 °C until LC-MS/MS analysis.

#### LC-MS/MS

GlyGly-K modified peptides isolated from B cells shown in Figure 3A were re-dissolved in 20 µL loading solvent A (0.1% TFA in water/ACN (98:2, v/v)) of which 10 µL were injected for LC-MS/MS analysis on an Ultimate 3000 RSLCnano system (ThermoFisher scientific) in-line connected to a Q Exactive HF mass spectrometer (Thermo). Trapping was performed at 10 μL/min for 4 min in loading solvent A on a 20 mm trapping column (100 μm internal diameter (I.D.), 5 μm beads, C18 Reprosil-HD, Dr. Maisch, Germany). Peptides were separated on a 200 cm long micro pillar array column (µPAC™, PharmaFluidics) with C18-endcapped functionality. This column consists of 300 µm wide channels that are filled with 5 µm porous-shell pillars at an inter pillar distance of 2.5 µm. With a depth of 20 µm, this column has a cross section equivalent to an 85 µm inner diameter capillary column. Peptides isolated from B cells were eluted from the analytical column by a non-linear gradient from 2 to 55% solvent B (0.1% FA in water/acetonitrile (2:8, v/v)) over 120 minutes at a constant flow rate of 300 nL/min, followed by 20 min in 99% solvent B. The column was then re-equilibrated with 98% solvent A for 10 min. For the analyses of peptides isolated from B cells, the column temperature was kept constant at 50 °C in a column oven (CoControl 3.3.05, Sonation). The mass spectrometer was operated in positive and data-dependent mode, automatically switching between MS and MS/MS acquisition for the 8 most abundant ion peaks per MS spectrum for the analyses of the peptides isolated from B cells. Full-scan MS spectra (375–1,500 m/z) were acquired at a resolution of 60,000 (at 200 m/z) in the orbitrap analyzer after accumulation to a target value of 3E6 for a maximum of 60 ms. The most intense ions above a threshold value of 8.3E3 were isolated in the trap with an isolation window of 1.5 Da for maximum 120 ms to a target AGC value of 1E5 for the analysis of peptides isolated from B cells. Only precursor ions with a charge state from 2 to 6 were selected for MS/MS. Peptide match was set on “preferred” and isotopes were excluded. Dynamic exclusion time was set to 12 s. Fragmentation was performed at a normalized collision energy of 28%. MS/MS spectra (200-2,000 m/z) were acquired at fixed first mass 145 m/z at a resolution of 15,000 (at 200 m/z) in the Orbitrap analyzer. MS/MS spectrum data type was set to centroid. The polydimethylcyclosiloxane background ion at 445.12003 was used for internal calibration (lock mass).

Dried peptides of the shotgun aliquots were re-dissolved in loading solvent A (0.1% TFA in water/ACN (98:2, v/v)) and 3 µg peptides of each sample was injected for LC-MS/MS analysis on an Ultimate 3000 RSLC nanoLC (Thermo Scientific, Bremen, Germany) in-line connected to an LTQ-Orbitrap Elite (Thermo Fisher Scientific, Bremen, Germany) equipped with a pneu-Nimbus dual ion source (Phoenix S&T). Trapping was performed at 10 μL/min for 4 min in solvent A on a 20 mm trapping column (made in-house, 100 μm internal diameter (I.D.), 5 μm beads, C18 Reprosil-HD, Dr. Maisch, Germany) and the sample was loaded on a reverse-phase column (made in-house, 75 µm I.D. x 250 mm length, 1.9 µm beads C18 Reprosil-HD, Dr. Maisch). Peptides were eluted by a non-linear increase from 1 to 56% MS solvent B (0.1% FA in water/ACN (2:8, v/v)) over 145 minutes, at a flow rate of 250 nL/min, followed by a 15-minutes wash reaching 99% MS solvent B and re-equilibration with MS solvent A (0.1% FA in water/ACN (2:8, v/v)). The mass spectrometer was operated in data-dependent, positive ionization mode, automatically switching between MS and MS/MS acquisition for the 20 most abundant peaks in a given MS spectrum. The source voltage was 3.1 kV, and the capillary temperature was 275°C. In the LTQ-Orbitrap Elite, full scan MS spectra were acquired in the Orbitrap (m/z 300−2,000, AGC target 3 × 10^6^ ions, maximum ion injection time 100 ms) with a resolution of 60 000 (at 400 m/z). The 20 most intense ions fulfilling predefined selection criteria (AGC target 5 × 10^3^ ions, maximum ion injection time 20 ms, spectrum data type: centroid, exclusion of unassigned and 1 positively charged precursors, dynamic exclusion time 20 s) were then isolated in the linear ion trap and fragmented in the high pressure cell of the ion trap. The CID collision energy was set to 35% and the polydimethylcyclosiloxane background ion at 445.120028 Da was used for internal calibration (lock mass).

For the pulldown-MS analyses shown in Figure 3E, dried peptides were re-dissolved in 20 µl loading solvent A of which 13 µL were injected for LC-MS/MS analysis as described for the analysis of peptides isolated from B cells. Peptides were eluted from the analytical column by a non-linear gradient from 2 to 56% solvent B over 150 minutes at a constant flow rate of 250 nL/min, followed by 10 min in 99% solvent B. The column was then re-equilibrated with 98% solvent A (0.1% FA in water) for 20 min. The column temperature was kept constant at 50 °C in a column oven (CoControl 3.3.05, Sonation). The mass spectrometer was operated as described above with minor modifications. It was run in positive and data-dependent mode, automatically switching between MS and MS/MS acquisition for the 16 most abundant ion peaks per MS spectrum. Full-scan MS spectra (375–1,500 m/z) were acquired at a resolution of 60,000 (at 200 m/z) in the orbitrap analyzer after accumulation to a target value of 3E6 for a maximum of 60 ms. The most intense ions above a threshold value of 1.3E4 were isolated in the trap with an isolation window of 1.5 Da for maximum 80 ms to a target AGC value of 1E5.

For the pull down-MS analyses shown in Figure 3H, dried peptides were re-dissolved in 20 µl loading solvent A of which 5 µL were injected for LC-MS/MS analysis as described above for the analysis of peptides isolated from B cells. Peptides were eluted from the analytical column as described for the analysis of peptides isolated from B cells, but over a gradient for 90 minutes. The column temperature was kept constant at 50 °C in a column oven (CoControl 3.3.05, Sonation). The mass spectrometer was operated as described above, but only the top 12 most abundant ion peaks were fragmented per MS spectrum. Fragmentation was performed as described for the analysis of peptides isolated from B cells but the dynamic exclusion time was set to 20 s.

#### LC-MS/MS data analysis

Data analysis was performed with MaxQuant (version 1.5.7.4, 1.5.8.3, 1.6.1.0 or 1.6.11.0) using the Andromeda search engine with default search settings including a false discovery rate set at 1% on both the peptide and protein level. The analysis of the B cells shown in Figure 3B was performed with two different searches to analyze the spectra generated from the analysis of the GlyGly-K enriched samples and the shotgun samples. Spectra from the pull down-MS shown in Figure 3E were searched twice: for the identification of GlyGly-K modified peptides and for the identification of proteins. Spectra from the pull down-MS shown in Figure 3H were searched once: for the identification of GlyGly-K modified peptides. Spectra were interrogated against proteins in the Uniprot/Swiss-Prot database ([www.uniprot.org]). The database released on January 2018 containing 16,946 mouse protein sequences were used for the analysis of Fig. 3B. The database released on September 2017 or January 2020 containing 20,226 or 20,595 human protein sequences was used for the pulldown-MS shown in Figure 3E and 3H, respectively. Mass tolerance for precursor ions was set to 4.5 ppm for all analyses. Mass tolerance for fragment ions was set to 20 ppm (orbitrap analyzer) for the analysis of the GlyGly-K modified peptides and the pulldown-MS, while it was set to 0.5 Da (ion trap analyzer) for the shotgun samples. Enzyme specificity was set as C-terminal to arginine and lysine (trypsin), also allowing cleavage at arginine/lysine–proline bonds with a maximum of three missed cleavages for the identification of GlyGly-K modified peptides or set to two missed cleavages for the identification of protein. Carbamidomethylation of cysteine residues was set as a fixed modification in the analyses of B cells. In all analyses, variable modifications were set to oxidation of methionine (to sulfoxides) and acetylation of protein N-termini. To identify GlyGly-K modified peptides, GlyGly modification of lysine residues was set as an additional variable modification. In the pulldown-MS, phosphorylation of Serine, Threonine or Tyrosine was set as an additional variable modification. Score for modified peptides was set to 40 or 30 for the identification of proteins or GlyGly-K modification sites, respectively. In the pulldown-MS shown in Figure 3E, score for modified peptides was set to 35 for the identification of proteins and GlyGly-K modification sites. In all searches, matching between runs was enabled with an alignment time window of 20 min and a matching time window of 1 min. Only proteins with at least one peptide were retained to compile a list of identified proteins. In the analysis of B cells, 3,273 mouse proteins, and 7,933 GG-modification sites were identified. In the pulldown-MS shown in Figure 3E, 1,389 human proteins and 126 GG-modification sites were identified. In the pulldown-MS shown in Figure 3H, 4,417 human proteins and 554 GlyGly-K modification sites were identified. Proteins were quantified by the MaxLFQ algorithm integrated in the MaxQuant software. A minimum of two ratio counts from at least one unique peptide was required for quantification in all analyses except for the pulldown-MS for which a minimum of two ratio counts from at least one unique or razor peptide was required.

Further data analysis was performed with the Perseus software (version 1.5.5.3, 1.6.1.1 or 1.6.2.3). For the analysis of the shotgun data, the proteinGroups table from MaxQuant was loaded in Perseus, reverse database hits were removed as well as proteins only identified by sites. The LFQ intensities were log2 transformed and replicate samples were grouped as above. Proteins with less than three valid values in at least one group were removed and missing values were imputed from a normal distribution around the detection limit. To reveal the proteins that were significantly regulated, sample groups were defined based on genotype (WT-USP8 vs. USP8_C748A_) and a t-test (FDR = 0.05 and S0 = 1) was performed to compare the intensity of the proteins between the two genotypes. Quantified proteins (n = 2,251) and the results of the t-tests are shown in Figure 3B. For the analysis of GlyGly-K modification sites, the GlyGly (K) Sites table from MaxQuant was loaded and reverse database hits were removed. The site table was expanded, intensities were log2 transformed and normalized for each sample by subtracting the median intensity. Replicate samples were grouped, sites with less than three valid values in at least one group were removed, and missing values were imputed from a normal distribution around the detection limit. To reveal sites that were significantly regulated, sample groups were defined based on genotype (WT-USP8 vs. USP8_C748A_) and a t-test (FDR = 0.05 and S0 = 1) was performed to compare the intensities of the sites between the two genotypes. Quantified GlyGly-K modified sites (n = 5,824) and the results of the t-tests are shown in Figure 3B. In all analyses, the same workflow as described above was applied for the analysis of GlyGly-K modification sites or for the analysis of proteins. For the pulldown-MS analysis shown in figure 3E, 845 proteins 49 GG-modification sites were evaluated. For the pulldown-MS analysis shown in Figure 3H, only identified GlyGly-K modification sites that are derived from NEDD8 or Ubiquitin proteins were displayed in the heatmap.

#### Statistical analysis

We ensured that our sample sizes matched those generally used in the field. T-testing, or 1-way or 2-way ANOVA and Tukey’s post hoc tests were applied and differences with a P value of <0.05 were considered significant. Excel and GraphPad Prism 10 were used for statistical testing if not mentioned otherwise.

The transcriptomic data and clinical data of the Multiple Myeloma Research Foundation (MMRF) CoMMpass study was retrieved from the Genomic Data Commons (GDC) data portal. Raw Counts were normalized for counts per million (CPM). The remaining samples were sorted according to the expression of USP8 and were stratified in two groups (<MEDIAN and > median). Kaplan-Meier overall survival plots on these three groups of samples were generated using the function TCGAanalyze_survival from the R package TCGAbiolinks v2.30[89].

The boxplot with patients stratified by International Staging System (ISS) was generated using ggplot2 v3.4.4 and ggpubr v0.6.

## Data availability

All MS proteomics data have been deposited to the ProteomeXchange Consortium via the PRIDE partner repository. The Shotgun analysis and the anlysis of GlyGly-K modification sites shown in Figure 3B were registered under the data set identifier PXD024395 (Username; Password: A6X1HcXc). The pull down-MS analysis shown in Figure 3E was registered under the data set identifier PXD024755 (Username; Password: Qgnx23Ev). The pull down-MS analysis shown in Figure 3H was registered under the data set identifier PXD024429 (Username; Password: FSW5Cl8e).

Single cell RNA-seq analysis data are available at https://www.ncbi.nlm.nih.gov/geo/query/acc.cgi?acc=GSE244334 (access token for reviewers: sxsdsecavdyxbqj).

## Supplemental Information

**Fig. S1.** Defect in B-cell development in *Usp8*^f/f^*Cd19*-Cre mice. (A) Marginal GADS expression in B cells as compared to T cells. Splenic T and B cells were enriched and stimulated for the times indicated with anti-CD3/CD28 or anti-IgM F(ab’) fragment antibody, respectively. Lysates were analyzed by WB. (B) Southern blot analysis of NcoI-digested genomic DNA from *Usp8*^f/f^ control and *Usp8*^f/f^*Cd19*-Cre B cells reveals efficient *Usp8* gene recombination. Deletion is indicated by a 6kb band representing the deleted allele. The 5kb band represents the floxed allele. (C) USP8 and actin expression in splenic B cells were analyzed by WB. (D) Flow cytometry analysis of B-cell subsets in spleen, LN, and BM of *Usp8*^+/+^*Cd19*-Cre mice and *Usp8*^f/f^*Cd19*-Cre mice. Pre-gating is indicated at the upper left of the FACS plots. Results are representative of 4 independent experiments. (E) Flow cytometry analysis of B1a- and B1b-cell subsets in spleen and peritoneal cavity (PCav), and B2 cell subsets in PCav of *Usp8*^+/+^*Cd19*-Cre and *Usp8*^f/f^*Cd19*-Cre mice. Pre-gating is indicated at the upper left of the FACS plots. The percentages and total numbers of B1 and B2 cell subsets are shown on the right (n=4, unpaired 2-tailed t-test, mean+/-sd). (F) Immunohistochemistry staining of paraffin-embedded spleen sections with hematoxylin/eosin (H&E), anti-B220, and anti-CD3 (brown) (scale bar 200 µm), and co-staining of frozen spleen sections as indicated (scale bar 100 µm). (G) Proliferation (BRDU incorporation) of purified splenic CD19^+^ *Usp8*^f/f^ and *Usp8*^f/f^*Cd19*-Cre B cells stimulated with anti-IgM in the absence or presence of anti-CD40, or with the TLR4 and TLR9 agonists “synthetic Monophosphoryl Lipid A” (MPLA) and CpG, respectively (paired, 2-tailed t-test representing n=3 sex-matched littermates, mean+/-sd). (H) Quantification of Ki67^+^ *Usp8*^f/f^ and *Usp8*^f/f^*Cd19*-Cre B cells in proliferating Ki67^+^ B-cell areas (n≥10, unpaired 2-tailed t-test, mean+/-sd). Scale bar, 100µm. (I) B-cell subsets in BM, spleen, LN, and PCav of *Usp8*^+/f^mb1-Cre mice and *Usp8*^f/f^*mb1*-Cre mice were analyzed by flow cytometry. Pre-gating is indicated at the upper left of the respective FACS plots. Results are representative of 3 independent experiments. Source data are provided as source data file.

**Fig. S2.**

Altered B-cell subsets, but normal myeloid and T-cell distribution, and inflammatory blood cytokine levels in B cell-specific Usp8 KO mice. (A) UMAP projection of B cell enriched splenocytes colored by graph-based clusters. (B) Flow cytometry analysis of splenic myeloid cells derived from *Usp8*^f/f^ and *Usp8*^f/f^*Cd19*-Cre mice (n=4, unpaired 2-tailed t-test, mean+/-sd). (C) Expression scores based on Mann-Whitney U statistic of 10 genes specific for the cell type reported[28] in UMAP projection. (D) Volcano plots of differential gene expression analysis of MZB, FoB, PC, and GC-like B cells. Red dots indicate significance for adjusted P value and log_2_ fold change. (E) Flow cytometry analysis of splenic Tcell subsets derived from *Usp8*^f/f^ and *Usp8*^f/f^*Cd19*-Cre mice. nT, naïve T cells; TEM, effector memory T cells; Treg, regulatory T cells (n=4, unpaired 2-tailed t-test, mean+-s/d). (F) Inflammatory cytokine levels in serum of *Usp8*^f/f^ and *Usp8*^f/f^*Cd19*-Cre mice (n=8, unpaired 2-tailed t-test, mean+-s/d). Source data are provided as source data file.

**Fig. S3.** *Cd19*-Cre-mediated deletion of USP8 alters splenic B-cell compartment signaling. (A) *Usp8*^f/f^ and *Usp8*^f/f^*Rosa26*-CreERT2 splenic B cells were expanded in medium containing IL-4 and CD40 for 24 h followed by OHT addition and further expansion for 48 h or 72 h. Lysates were subjected to WB analysis. Results are representative of at least two independent experiments. (B) Gene expression of USP8, Roquin and Malt1 genes in splenic *Usp8*^f/f^ and *Usp8*^f/f^*Cd19*-Cre B-cell subsets. Black dots represent a single cell each. (C) Ca^2+^ influx in anti-IgM-stimulated B cells. Splenic *Usp8*^f/f^ and *Usp8*^f/f^*Cd19*-Cre B cells were purified with biotinylated anti-CD19 antibodies and IMag Streptavidin Particles plus-DM and starved for 2 hours. B cells were loaded with indo-1-AM (5μg/ml) and activated with anti-IgM F(ab’) fragment antibody (10μg/ml). The graph shows the ratio of bound to unbound indo-1-AM as a measure of Ca^2+^ influx (n=3). (D) Splenic B cells were purified, starved for 2 h, stimulated for the indicated times with anti-IgM F(ab’) fragment antibody, and analyzed by WB. (E) *Usp8*^f/f^ and *Usp8*^f/f^*Rosa26*-CreERT2 splenic B cells were expanded in medium containing IL-4 and CD40 for 24 h followed by OHT addition and further expansion for 48 h. Cells were stimulated as in (D) and lysates were subjected to WB analysis. Results in D and E are representative of at least two independent experiments. (F) Cells were expanded as in (E) for 72 h after OHT addition and Ca^2+^ influx was analyzed as in (C) (n=3). (G) Upon oral gavage of tamoxifen, Roquin1/2 depleted *Rc3h1/2*^f/f^ *Rosa26-*CreERT2 B cells and *Rosa26-*CreERT2 control cells were enriched from spleens and Ca^2+^ influx was analyzed as in (C) (n=4). (H) Pathway enrichment analysis on genes upregulated in B-cell subsets of the indicated genotype. Source data are provided as source data file.

**Fig. S4.** Role of K11- and K48-linked NEDD8 modifications in the formation of mixed ubiquitin/NEDD8 chains induced by catalytically inactive USP8. (A) HEK293T cells were transfected with WT-USP8-FLAG or USP8(C786A)-FLAG in the presence of the indicated HA-NEDD8 variants and lysates were analyzed by WB. (B) Selected samples shown in (A) were analyzed by WB as indicated. Results are representative of 2 independent experiments. Source data are provided as source data file.

**Fig. S5.** (A) Viability assay of BTZ resistant KMS-18-T21A MM cells upon treatment with 10nM (left panel) or 25 nM (right panel) BTZ in the presence of increasing DUB-IN-2 concentrations (n=3, mean+/-sd). (B) Cell viability of KMS-27 WT and BTZ-resistant mutant MM cells (A49V) treated with CFZ (3 nM), DUB-IN-2 (25 µM), or a combination of both for the times indicated (n=3, 2-way ANOVA, Tukey’s post hoc test, mean+/-sd). (C) WB analysis of KMS-27/KMS-27-A49V MM cells after exposure to CFZ and/or DUB-IN-2 for 6 h. Quantification of ubiquitin modification shown on top the blot represents the average of 2 independent experiments normalized to actin and DMSO controls. Source data are provided as source data file. (D) BM cells from selected MM patients who either insufficiently responded to BTZ-containing induction therapy or lost their response during BTZ-containing treatment (defined as “BTZ-resistant”, n=4) and patients responding well to BTZ-containing induction therapy (defined as “BTZ non-resistant”, n=4) were incubated with solvent (DMSO), or increasing concentrations of BTZ, or DUB-IN-2 for 24 hours and viability was measured by AnnexinV/7AAD staining. Unpaired 2-tailed t-test, mean+/- sd. (E) KMS-27-A49V cells transfected with pooled NT siRNA or USP8 siRNA_2 for 96 h, and NT siRNA transfected cells treated with BTZ (25 nM) and/or DUB-IN-2 (25µM) for 24 h were analyzed by AO staining. Results are representative of 5 pictures taken from 2 independent experiments, respectively.

**Fig. S6.** (A) Cell viability of KMS-18-T21A MM cells treated with the indicated concentrations of OTUB1/USP8 inhibitor 61 (OUI61) in the absence or presence of BTZ (10 nM) (n=3, 2-way ANOVA, Tukey’s post hoc test, mean+/-sd). (B) OUI61 exerts a caspase 3-independent toxic effect on cell viability. KMS-18-T21A cells were treated with the indicated concentrations of OUI61 for 24 h and subsequently analyzed by WB. (C) Proteasome activity of KMS-18 MM cell lysates treated with MG132 (positive control), DUB-IN-2 (12.5 µM), and/or BTZ (7.5 nM) was assessed using fluorogenic substrates specific for chymotrypsin-like (Suc-LLVY-AMC), trypsin-like (Boc-LRR-AMC), and caspase-like (Z-LLE-AMC) activities. (D) Lack of BM infiltration of KMS-18/KMS-18-T21A MM cells in NSG mice that had been treated with either control (0.9NaCl/PEG300 1:1 (v/v)), BTZ (0.6mg/kg, twice a week), or BTZ combined with DUB-IN-2 (n=6) using 5mg/kg 5 times a week over a period of 13 days, followed by 2mg/kg twice a week (mean+/-sd).

**Fig. S7.** (A) Kaplan Meier survival analysis of MM patients included in the MMRF CoMMpass study according to USP8 expression at time of diagnosis (Log rank test). (B) Box plot of USP8 expression in patients of different MM disease stages (Kruskal-Wallis test).

