## Supplementary figures and images for "USP8 Controls Proteostasis Pathways in B Cells and Multiple Myeloma"

### Suppl Figure 1

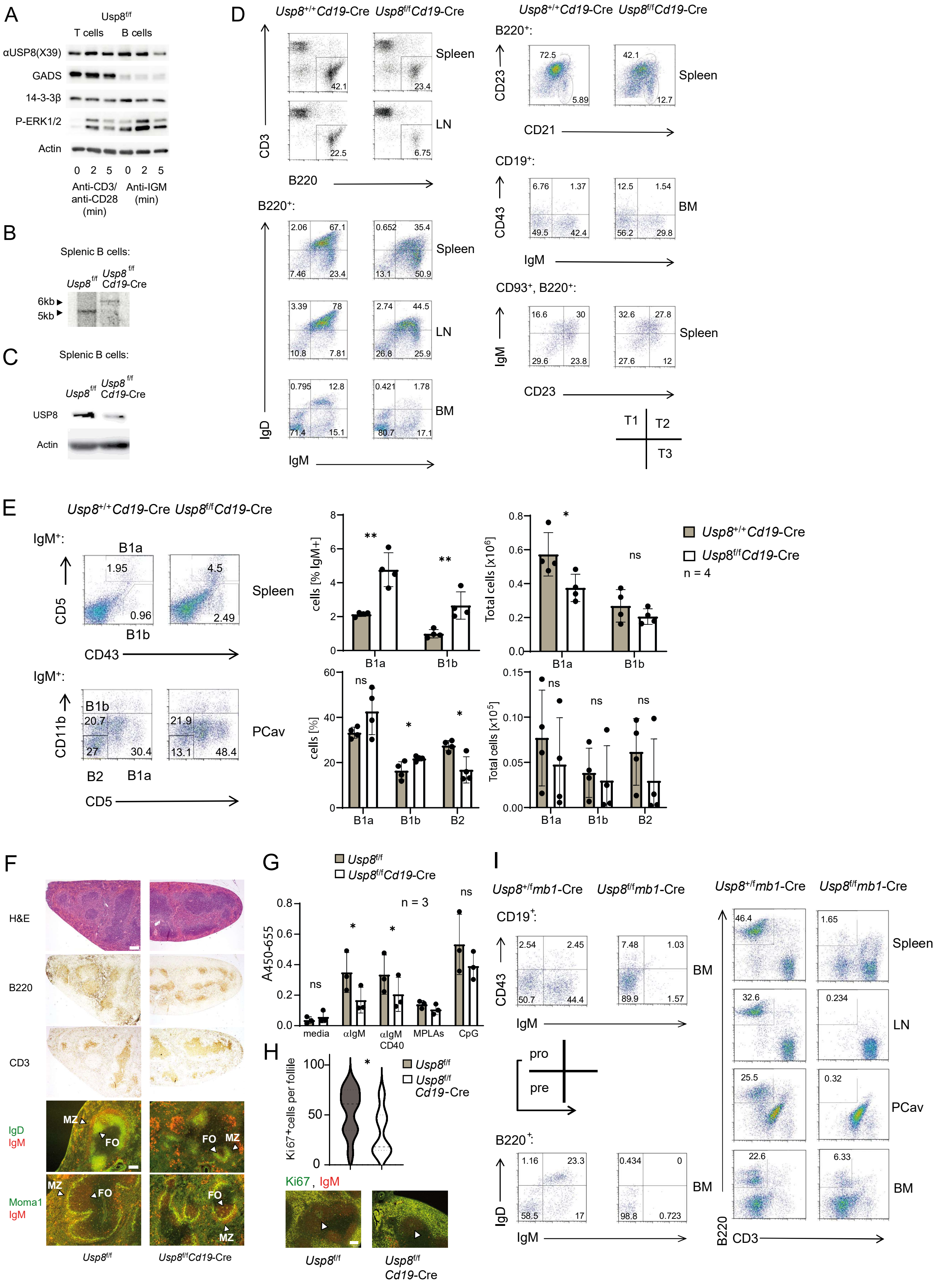


Dufner et al., Figure S1

### Suppl Figure 2

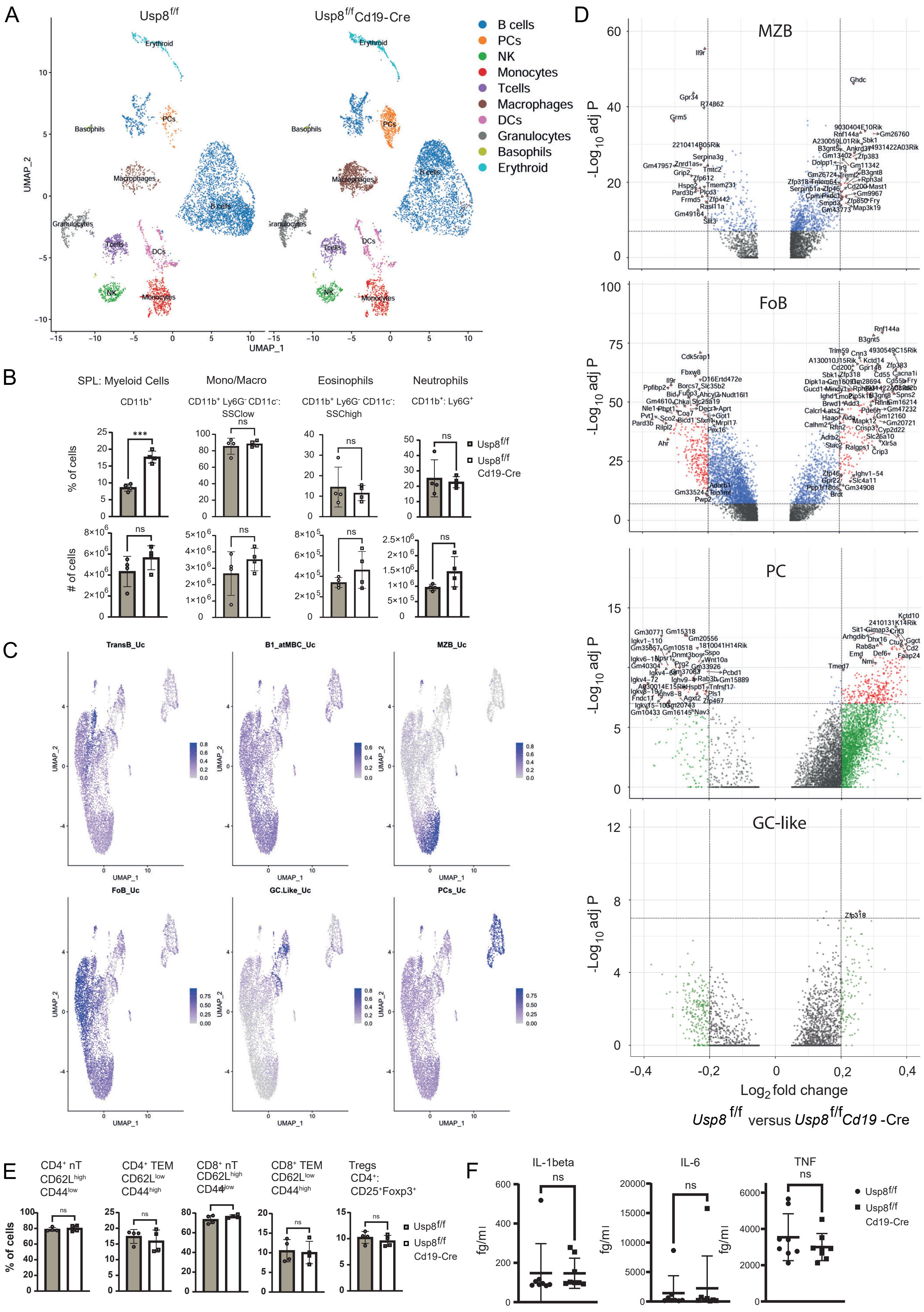


Dufner et al., Figure S2

### Suppl Figure 3

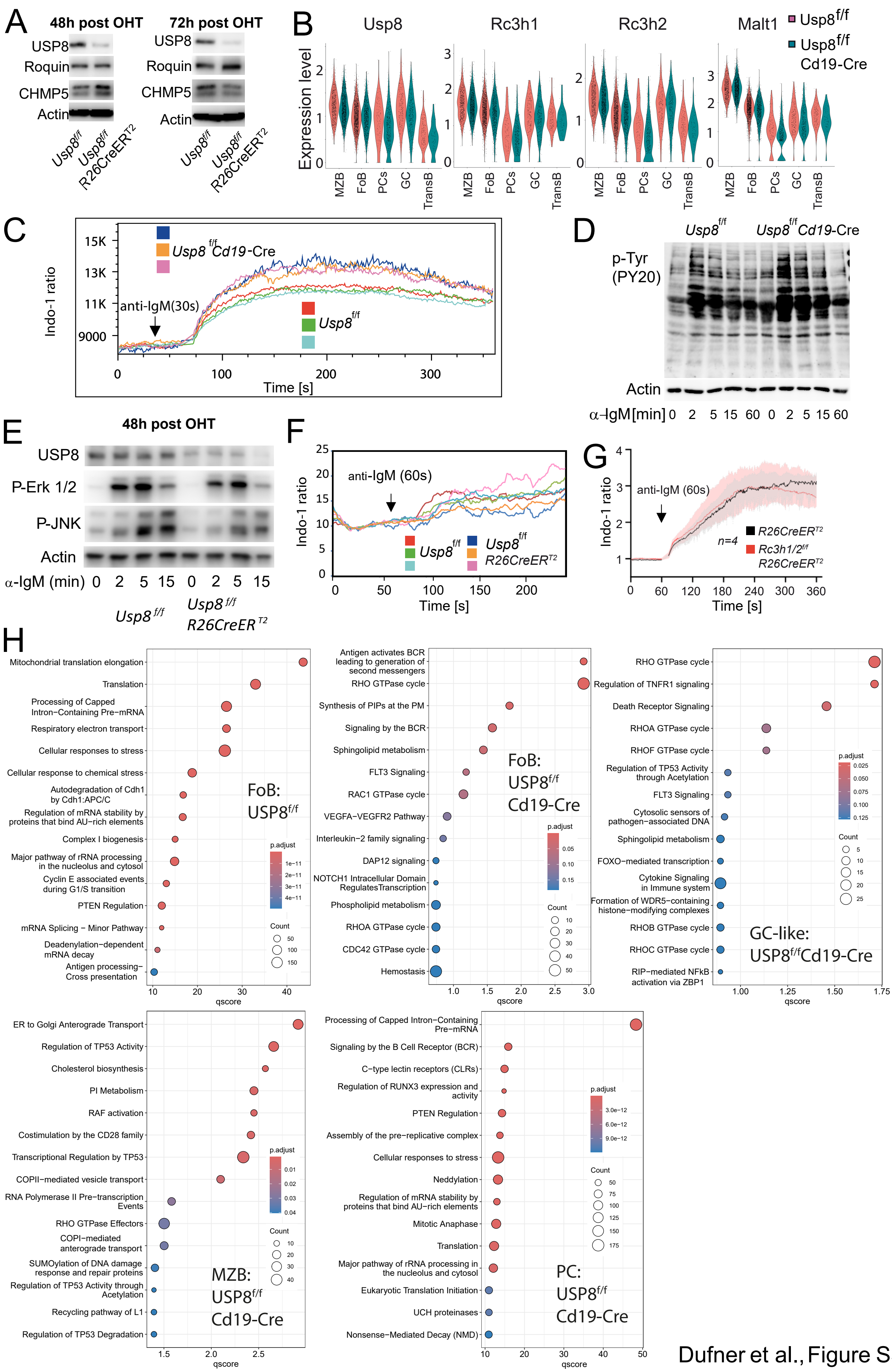


Dufner et al., Figure S3

### Suppl Figure 4

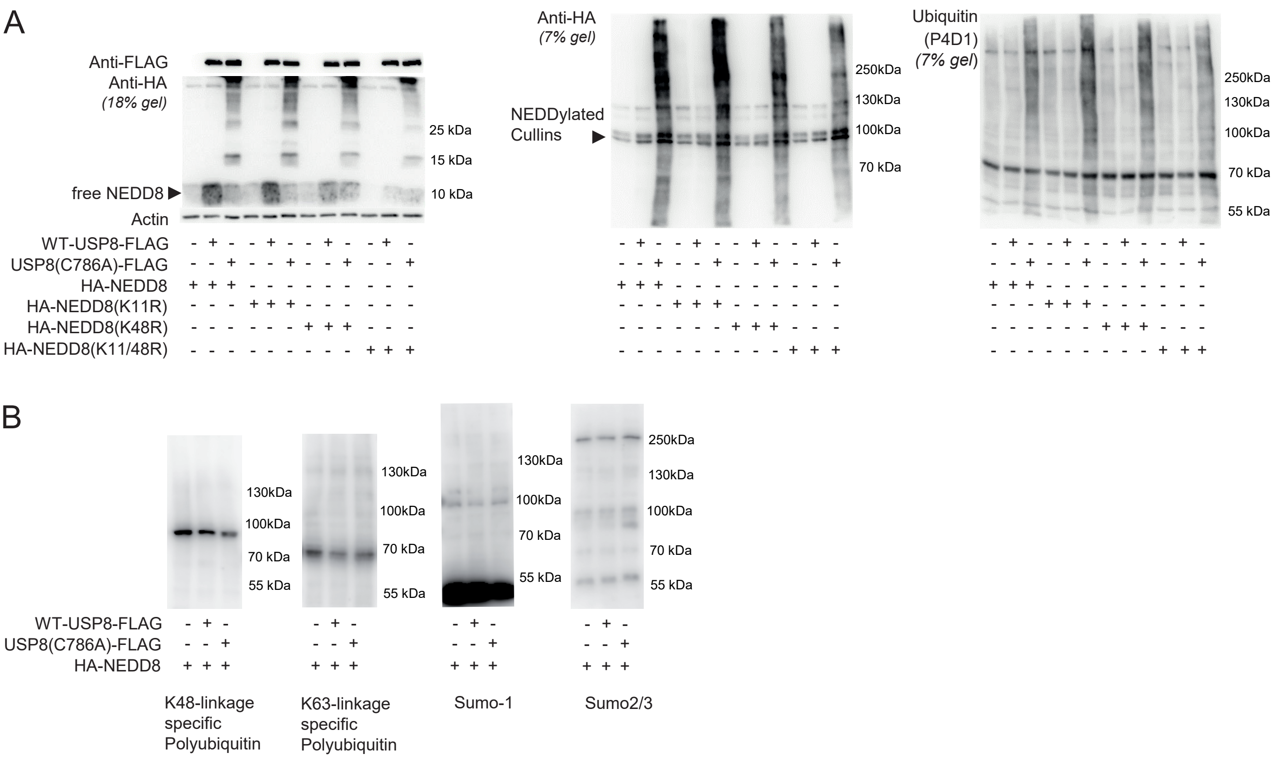


Dufner et al. Figure S4

### Suppl Figure 5

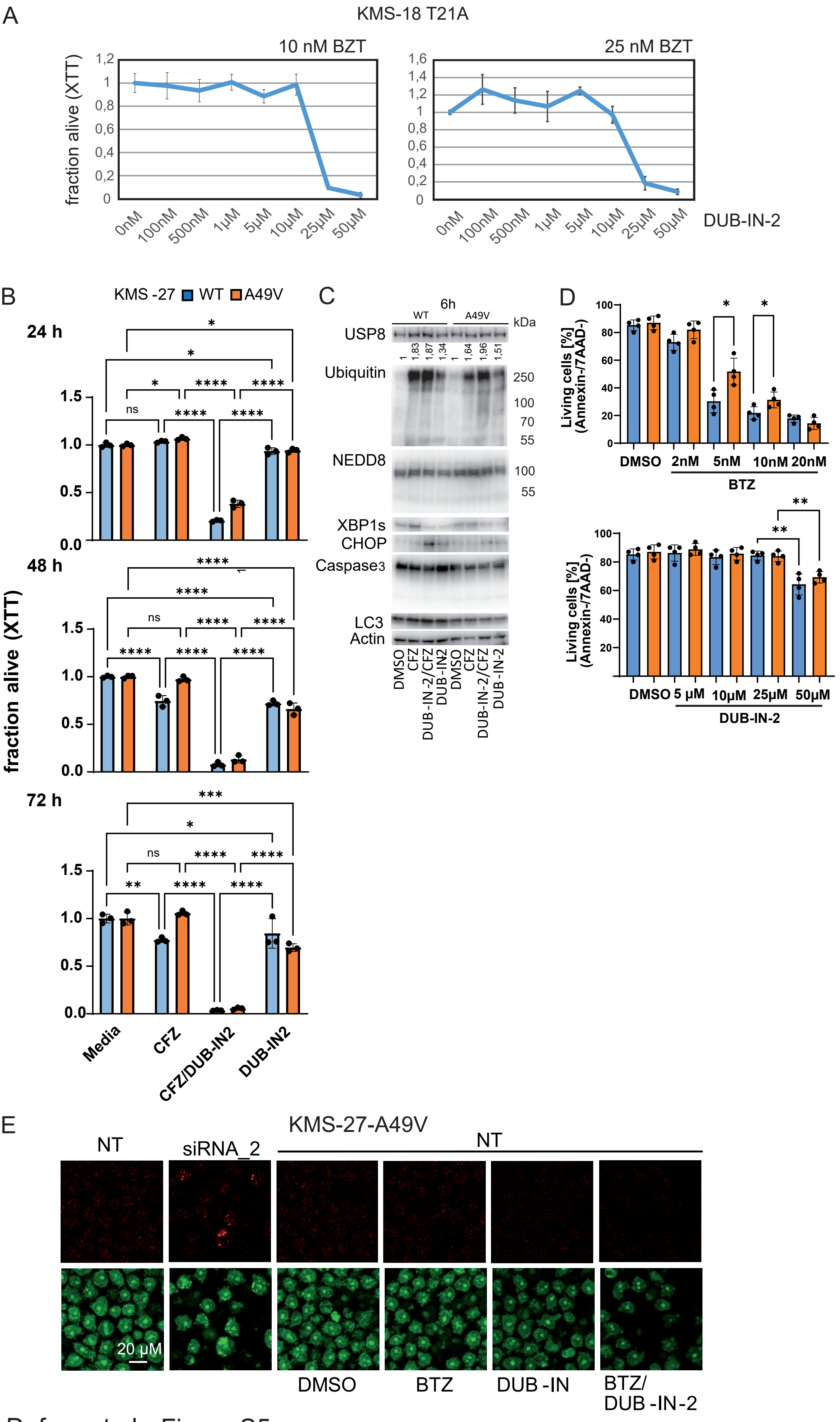


Dufner et al., Figure S5

### Suppl Figure 6

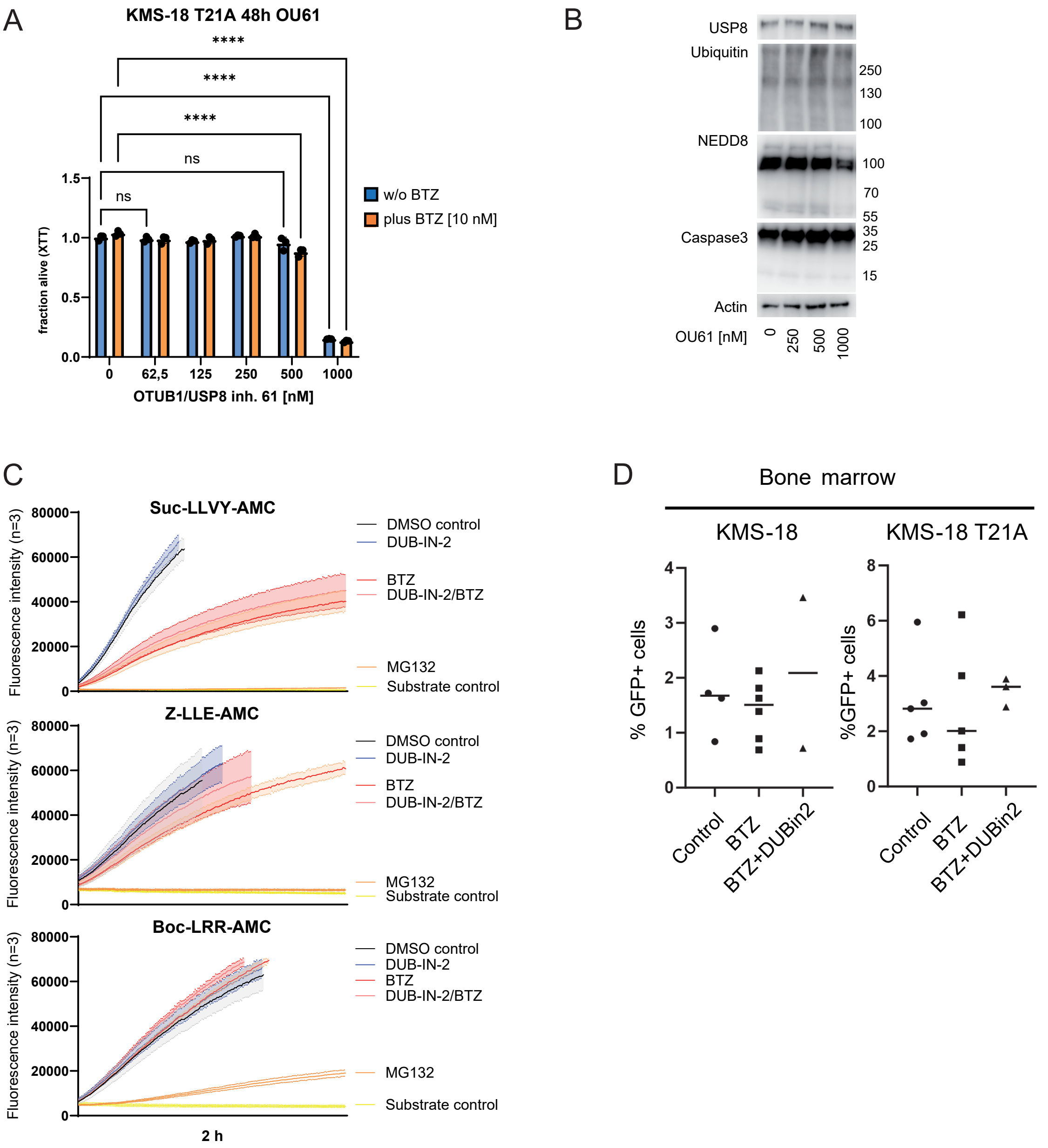

### Suppl Figure 7

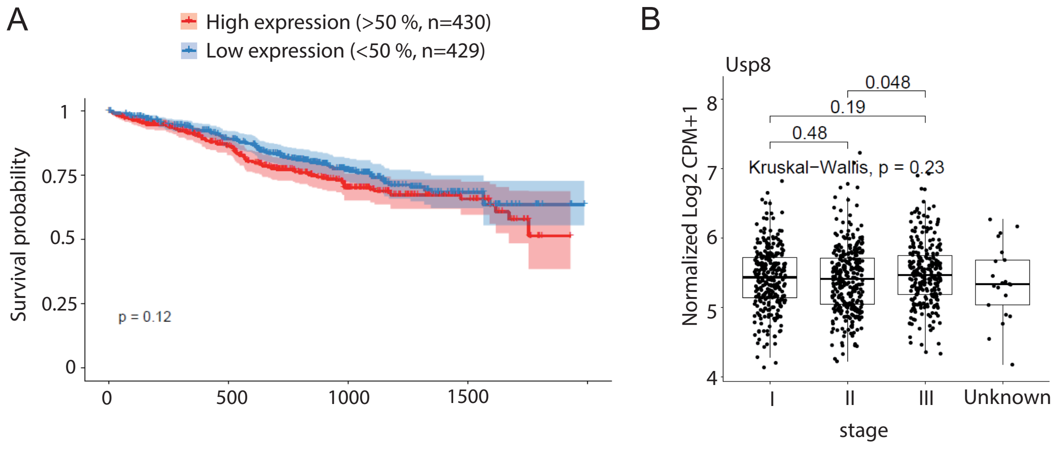


Dufner et al., Fig. S7
