## Supplementary material for "USP8 Controls Proteostasis Pathways in B Cells and Multiple Myeloma": Table S1

**Table S1.** Neddylation-related genes upregulated in *Usp8*^f/f^*Cd19*-Cre PCs.

| **Commd8** | **Psmb1** | **Psma1** | **Psmc2** | **Uba3** |
| --- | --- | --- | --- | --- |
| **Ube2d3** | **Psme2** | **Nedd8** | **Dda1** | **Klhl42** |
| **Psmb8** | **Ube2d2a** | **Fem1a** | **Psmb7** | **Rbbp7** |
| **Fbxl8** | **Psmd13** | **Keap1** | **Ercc8** | **Btbd1** |
| **Psmd4** | **Lrrc41** | **Psma4** | **Dcun1d1** | **Fbxo22** |
| **Psmb9** | **Psmc5** | **Uba52** | **Gan** | **Cul3** |
| **Asb13** | **Psmf1** | **Uchl3** | **Asb6** | **Fbxo7** |
| **Psmd5** | **Psma2** | **Cops5** | **Psma6** | **Fbxl14** |
| **Psme2b** | **Nub1** | **Commd6** | **Psmb4** | **Tulp4** |
| **Ufd1** | **Wsb2** | **Psme1** | **Wsb1** | **Nploc4** |
| **Dcaf11** | **Dcaf17** | **Ube2d1** | **Psmd12** | **Commd10** |
| **Cops7b** | **Psma3** | **Dcaf7** | **Commd9** | **Psmd6** |
| **Psmd10** | **Cul4b** | **Psmc1** | **Hif1a** | **Fbxw4** |
| **Psmd14** | **Dcaf13** | **Cand1** | **Mul1** | **Klhl20** |
| **Dcun1d5** | **Commd4** | **Ube2m** | **Ddb2** | **Ubxn7** |
| **Dcun1d2** | **Psmc3** | **Psma5** | **Spsb3** | **Psmd7** |
| **Psmb10** | **Psmd2** | **Ddb1** | **Cops4** | **Psmd11** |
| **Ube2f** | **Fbxo21** | **Commd2** | **Psmd1** | **Cul2** |
| **Fem1b** | **Psmc4** | **Cul5** | **Psmb2** | **Fbxo4** |
| **Psmc6** | **Fbxw9** | **Psma7** | **Cops2** | **Rnf7** |
| **Psmd8** | **Rbbp5** | **Commd7** | **Eloc** | **Rps27a** |
