## Supplementary material for "USP8 Controls Proteostasis Pathways in B Cells and Multiple Myeloma": Table S6

**Table S6.** Clinical characteristics of MM patients from whom samples were derived.

| **patient ID** | **bortezomib resistant/ refractory*** | **gender** | **age** | **BM sampling** | **number of prior therapy lines** | **prior therapy regimens**** | **details** |
| --- | --- | --- | --- | --- | --- | --- | --- |
| MM_01 | yes | female | 40 | at initial diagnosis | 0 | n.a. | insufficient response upon VCD induction therapy, switch to VRD/ Dara-Rd |
| MM_02 | yes | male | 62 | at initial diagnosis | 0 | n.a. | insufficient response upon VCD induction therapy, switch to Dara-VTD |
| MM_03 | yes | female | 53 | at initial diagnosis | 0 | n.a. | insufficient response upon BAD induction therapy, switch to Dara-Rd |
| MM_04 | yes | male | 67 | at initial diagnosis | 0 | n.a. | insufficient response upon BAD induction therapy, switch to Dara-Rd |
| MM_05 | yes | male | 82 | at initial diagnosis | 0 | n.a. | progressive disease upon 4 cycles of VD |
| MM_06 | yes | male | 52 | during treatment | 1 | VRD | insufficient response upon VCD induction therapy, switch to VRD |
| MM_07 | yes | female | 50 | upon alloSCT | 2 | VCD + autoSCT; Dara-VD + alloSCT | recurrent myeloma d+315 upon alloSCT, received prior bortezomib-containing treatment regimens |
| MM_08 | yes | female | 66 | upon alloSCT | 2 | VCD + autoSCT; Len/Pom-AD + alloSCT | myeloma d+45 upon alloSCT, received prior bortezomib-containing treatment regimens |
| MM_09 | yes | male | 67 | during treatment | 6 | Z-Dex + autoSCT; VD; Rd + alloSCT; Dara-Rd; KD; Isa-Pom-Dex | refractory myeloma upon 30 cycles of Isa-Pom-Dex, received prior bortezomib-containing treatment regimens |
| MM_10 | no | female | 71 | at initial diagnosis | 0 | n.a. | achieved PR before auto-SCT upon 4 cycles VCD and mobilization chemotherapy with CE |
| MM_11 | no | male | 72 | at initial diagnosis | 0 | n.a. | achieved PR before auto-SCT upon 4 cycles VCD and mobilization chemotherapy with CE |
| MM_12 | no | male | 66 | at initial diagnosis | 0 | n.a. | achieved PR before auto-SCT upon 4 cycles VCD and mobilization chemotherapy with CE |
| MM_13 | no | male | 62 | at initial diagnosis | 0 | n.a. | achieved PR before auto-SCT upon 4 cycles VCD and mobilization chemotherapy with IEV |
| MM_14 | no | male | 74 | at initial diagnosis | 0 | n.a. | achieved a VGPR upon 6 cycles VCD, subsequently de-escalation to VD |
| MM_15 | no | female | 63 | at initial diagnosis | 0 | n.a. | achieved a VGPR upon 7 cycles VCD/BAD, subsequently de-escalation to VD |

** Patients were either treatment-naive and had an insufficient response to bortezomib-containing induction treatment regimens (resistant) or their myeloma progressed upon bortezomib-containing treatment regimens (refractory)*

*** Treatment regimens: VCD – Bortezomib/ Cyclophosphamide/ Dexamethasone; BAD - Bortezomib/ Adriamycin/ Dexamethasone; Z-Dex – Idarubicin/ Dexamethasone; VRD – Bortezomib/ Lenalidomide/ Dexamethasone; (Dara)-Rd – (Daratumumab)/ Lenalidomide/ Dexamethasone; Dara-VTD – Daratumumab/ Bortezomib/ Thalidomide/ Dexamethasone; (Dara)-VD – Daratumumab/ Bortezomib/ Dexamethasone; Len/Pom-AD – Lenalidomide/ Pomalidomide/ Adriamycin/ Dexamethasone; KD – Carfilzomib/ Dexamethasone; Isa-Pom-Dex – Isatuximab/ Pomalidomide/ Dexamethasone; CE – Cyclophosphamide/ Etoposide; IEV –Iifosfamide/ Epirubicin/ Etoposide; autoSCT – autologous stem cell transplantation; alloSCT – allogenic stem cell transplantation*
