## Supplementary material for "USP8 Controls Proteostasis Pathways in B Cells and Multiple Myeloma": Table S7

**Bliss Score (CI) Table**

|  | KMS-18 | KMS-18T21A | KM-S27 | KMS-27A49V |
| --- | --- | --- | --- | --- |
| 24 h | 0,99396991 | 0,71384125 | 0,99757824 | 0,95268967 |
| 48 h | 0,98379361 | 0,65171124 | 0,94665343 | 0,95085692 |

CI = (E_A_+E_B_-E_A_*E_B_)/E_AB_

E_A_= Effect of drug A

E_B_=Effect of drug B

E_AB_=Effect of combination of Drug A and B

(0,9<CI<1,1 is considered additive; 0,7<CI<0,9 is considered mildly synergistic)
