## Supplementary material for "USP8 Controls Proteostasis Pathways in B Cells and Multiple Myeloma": Table S8

**Table S8.** Antibodies/target-specific binding molecule list.

| **Experiment** | **Marker/target-specific binding molecule** | **Clone ID** | **Source** |
| --- | --- | --- | --- |
| **Histology** | B220 | RA3-6B2 | BD Pharmingen |
|  | CD3 | CD3-12 | Serotec |
|  | IgD (biotinylated) | 11.26.c.2a | BioLegend |
|  | IgM | Polyclonal | Chemicon |
|  | Moma1 | MCA947 | AbD Serotec |
|  | Biotinylated peanut agglutinin (PNA) | # B-1075 | Vector Laboratories |
|  | Ki67 | TEC-3 | DAKO |
| **FACS** | Anti-CD16/CD32  (Fc Block^TM^) | 2.4G2 | BD Pharmingen |
|  | CD3ε | 500A2 | eBioscience |
|  | CD19 | MB19-1 | eBioscience |
|  | B220 | RA3-6B2 | BD Pharmingen |
|  | IgD | 11.26.c.2a | BD Pharmingen |
|  | IgM | II/41 | eBioscience |
|  | CD5 | 53-7.3 | BD Pharmingen |
|  | CD21 | 7G6 | BD Pharmingen |
|  | CD23 | B3B4 | BD Pharmingen |
|  | CD43 | S7 | BD Pharmingen |
|  | CD93 | AA4.1 | eBioscience |
|  | Ki67 | B56 | BD Pharmingen |
|  | T- and B-cell activation antigen | GL7 | BD Pharmingen |
|  | Fas | Jo2 | BD Pharmingen |
|  | CD138 | 281-2 | BioLegend |
|  | IRF4 | REA201 | Milteny |
|  | IgG1 | RMG1-1 | BioLegend |
|  | Blimp1 | 5E7 | BioLegend |
|  | IgM | RMM-1 | BioLegend |
|  | CD16/32 | 93 | BioLegend |
|  | CD19 | 6D5 | BioLegend |
|  | CD21/CD35 | 7E9 | BioLegend |
|  | CD25 | 3C7 | BioLegend |
|  | CD3 | 17A2 | BioLegend |
|  | CD335 | 29A1.4 | BioLegend |
|  | CD4 | GK1.5 | BD Pharmingen |
|  | CD44 | IM7 | BioLegend |
|  | CD62L | MEL-14 | BioLegend |
|  | CD8a | 53-6.7 | BioLegend |
|  | F4/80 | BM8 | BioLegend |
|  | FOXP3 | 150D | BioLegend |
|  | I-A/I-E | M5/114.15.2 | BioLegend |
|  | IgM | RMM-1 | BioLegend |
|  | Ly-6C | HK1.4 | BioLegend |
|  | Ly6G | 1A8 | BD Pharmingen |
|  | TCRbeta | H57-597 | BioLegend |
| **WB** | USP8(13aa) | Polyclonal | ^37^ |
|  | USP8 (X39) | Polyclonal | ^87^ |
|  | α,β,γ Actin | I-19 | Santa Cruz Biotechnology |
|  | FLAG | M2 | Sigma |
|  | HA | 3F10 | Roche |
|  | Ubiquitin | P4D1 | Santa Cruz Biotechnology |
|  | NEDD8 | Polyclonal, #2745 | Cell Signaling Technology |
|  | NEDD8 | Y297 | Abcam |
|  | phospho-p44/42 | 197G2 | Cell Signaling Technology |
|  | phospho-JNK | 81E11 | Cell Signaling Technology |
|  | phospho-p38 | 3D7 | Cell Signaling Technology |
|  | phospho-Akt | Polyclonal, #9271 | Cell Signaling Technology |
|  | phospho-IκBα | 14D4 | Cell Signaling Technology |
|  | Roquin1/2 | 3F12 | ^76^ |
|  | IκBα | Polyclonal, #9242 | Cell Signaling Technology |
|  | phosphotyrosine | PY20 | BD Biosciences |
|  | DR5 | E9D7D | Cell Signaling Technology |
|  | CHOP | L63F7 | Cell Signaling Technology |
|  | LC3A/B | Polyclonal, #4108 | Cell Signaling Technology |
|  | Caspase3 | Polyclonal, #9662 | Cell Signaling Technology |
|  | Caspase8 | D35G2 | Cell Signaling Technology |
|  | Caspase9 | C9 | Cell Signaling Technology |
|  | XBP1s | E9V3E | Cell Signaling Technology |
|  | SUMO-1 | Polyclonal, #4930 | Cell Signaling Technology |
|  | SUMO-2/3 | 18H8 | Cell Signaling Technology |
|  | Cullin-1 | EPR3103Y | Abcam |
|  | Cullin-3 | Polyclonal, #11107-1-AP | Proteintech |
|  | Cullin-4b | Polyclonal, #12916-1-AP | Proteintech |
|  | K48-linkage specific Polyubiquitin | Polyclonal, #4289 | Cell Signaling Technology |
|  | K63-linkage specific Polyubiquitin | D7A11 | Cell Signaling Technology |
